# Scaling perturbations: beyond genome-scale CRISPR screens

**DOI:** 10.64898/2026.01.16.699948

**Authors:** Anran Tang, Rico C. Ardy, Rafaela E. Mendes, Thomas M. Norman

## Abstract

CRISPR screens have become essential tools for systematically probing gene function from basic biology to drug discovery, yet important frontiers remain beyond genome scale. Probing regulatory elements, interpreting genetic variants, and mapping genetic interactions all challenge the sensitivity and scalability of existing approaches. Here we introduce two synergistic technologies to address these limitations. PORTAL (Perturbation Output via Reporter Transcriptional Activity in Lineages) shifts pooled genetics toward quantitative RNA phenotypes, encoding perturbation effects in expressed transcripts to enable single-molecule measurements with lineage or single-cell resolution. CAP cloning (Covalently closed Assembly Products) bypasses bacterial transformation to enable construction of ultrahigh-complexity lentiviral libraries. Combining these advances, we construct a genetic interaction map spanning 665,856 pairwise perturbations across 46 million clonal lineages—the largest exhaustive map in human cells and the first at this scale using a non-fitness phenotype. More broadly, this work charts a path toward comprehensive genetic interaction mapping in human cells.

## Introduction

Pooled CRISPR screens have transformed how genetics is performed in mammalian systems^1–6^. In one sense, the genome has been conquered: screens have been performed in thousands of cell lines^7,8^, genome-wide sgRNA libraries have gone through multiple iterations of improvement^9–15^, and a mature ecosystem of analytical software now exists^16–19^. However, important frontiers remain beyond genome scale. Probing regulatory elements^20–26^, understanding genetic variants^27–33^, and mapping genetic interactions^13,34–41^ all challenge the scalability and sensitivity of existing approaches.

A useful way to frame these challenges is to consider a constraint faced by almost all pooled screens: because they are ultimately read out using a fixed sequencing budget, there is a fundamental tradeoff between the number of cells that can be assayed and the amount of information obtained per cell. Different screening methods fall at different points along this continuum. A fitness screen is effectively a census, with each cell contributing a single count corresponding to the sgRNA it carries. Single-cell methods such as Perturb-seq^42,43^ or CROP-seq^44^ sit at the opposite extreme, extracting rich phenotypes from each cell in the form of a whole transcriptome. This richness, however, comes at an increased sequencing cost, limiting the number of perturbations that can be profiled. Reporter-based approaches, including fluorescence in situ hybridization (FISH) quantification of specific transcripts^45,46^ and transcriptional reporter constructs^47–49^, offer an appealing middle ground by converting perturbation effects into a targeted, quantitative molecular readout that can be measured efficiently at scale. In practice, however, existing implementations are constrained in throughput by reliance on cell sorting or by a need for complex cell-line engineering to introduce genomic landing pads that is impractical in many experimental settings.

Scaling the number of perturbations also leads to additional bottlenecks in steps that are often taken for granted. Library construction using standard cloning methods relies on bacterial transformation and propagation, which impose limits on library size and uniformity that can distort representation at high complexity, particularly when library elements have phenotypic effects in bacteria. As computational approaches^50^ expand the space of possible perturbations beyond the genes in the genome, this mismatch between what can be designed and what can easily be cloned is likely to become increasingly severe.

Here we introduce two synergistic technologies that address these constraints to enable a new scale of CRISPR screens. PORTAL (Perturbation Output via Reporter Transcriptional Activity in Lineages) rethinks pooled CRISPR screening by bringing key innovations from single-cell functional genomics back into the bulk screening context. PORTAL builds on the robustness of conventional lentiviral screens, but encodes all information in expressed transcripts rather than genomic DNA. This design enables incorporation of features such as unique molecular identifiers (UMI)^51^ for single-molecule quantification, clonal barcodes for lineage-resolved screens^52,53^, and combinatorial indexing^54^ to recover single-cell resolution when desired. To further enable experiments at unprecedented scale, we pair PORTAL with CAP cloning (Covalently closed Assembly Products). CAP cloning bypasses bacterial transformation by synthesizing and amplifying linear DNA molecules that are directly competent for lentiviral packaging, enabling straightforward construction of ultrahigh-complexity lentiviral libraries.

We apply these technologies to construct a genetic interaction (GI) map spanning 665,856 pairwise perturbations targeting 612 genes and measured across 46 million clonal lineages—the largest exhaustive interaction map measured to date in human cells, and the first at this scale using a non-fitness phenotype. Using AP-1 transcriptional activity as a broadly responsive readout, we recover interpretable structure across multiple levels of biological organization, including pathway-level relationships, subcomplex architecture within multi-protein machines, and functional connections between individual genes. Together, these results establish a strategy for functional genomics beyond genome scale, including comprehensive GI mapping across the human genome.

### Quantitative, lineage-resolved CRISPR screens using PORTAL

Traditional CRISPR screens read out perturbation identity by sequencing sgRNAs from genomic DNA, meaning each cell contributes at most a single molecule to the assay. Inspired by CiBER-seq^55^, CROP-seq^44^, and other single-cell functional genomics approaches, we developed PORTAL, a reporter-based screening method that instead reads out perturbation effects via RNA. By encoding perturbation identity and lineage information in expressed transcripts, PORTAL enables quantitative measurement of transcriptional phenotypes with single-molecule and clonal resolution. Because each clonal lineage provides an independent replicate measurement, this approach enables either reduced cell numbers per perturbation or dramatically increased experimental scale. As a practical benefit, RNA extraction is also simpler than genomic DNA extraction.

Our lentiviral vector for PORTAL encodes an sgRNA that specifies the genetic perturbation, a clonal barcode^52,53^, and two embedded Pol II transcripts within a single integrated cassette (Fig. 1A). A reporter transcript measures activity from an arbitrary perturbation-responsive promoter, while a nested constitutive promoter expresses an identity transcript that ensures each cell is detected regardless of reporter activity. The 3′ LTR contains the sgRNA expression cassette and clonal barcode, which together define clonal lineages arising from the same integration event. As a result, both reporter and identity transcripts carry perturbation and lineage information. Reporter expression is coupled to mCherry and identity expression to GFP (or puromycin), enabling orthogonal readout by flow cytometry.

**Figure 1.**
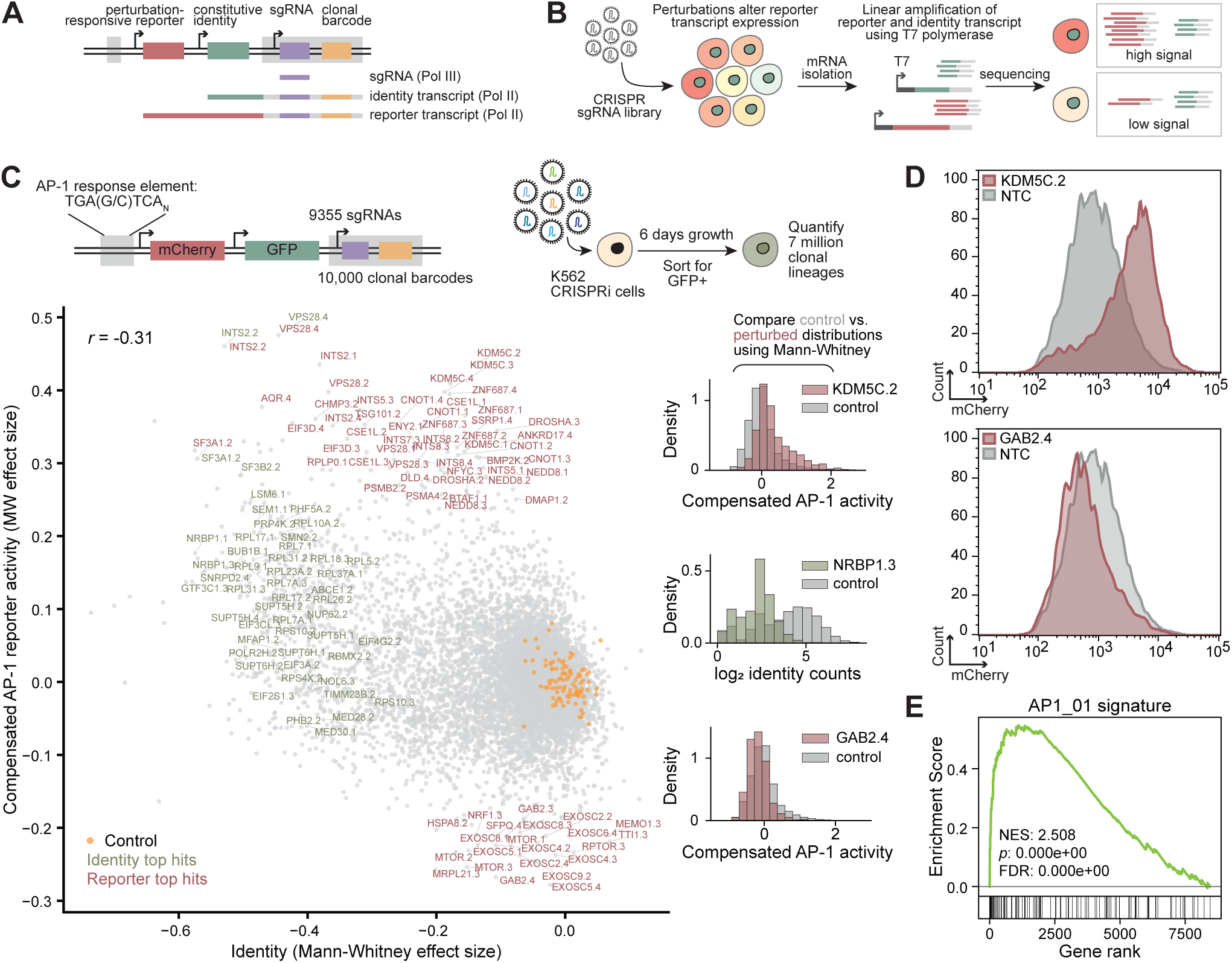
Development of PORTAL approach for lineage-resolved CRISPR screens. **A**, Schematic of the PORTAL lentiviral vector. A single integration encodes an sgRNA, a clonal barcode, and two Pol II transcripts: a reporter transcript driven by a perturbation-responsive promoter and an identity transcript driven by a nested constitutive promoter. The sgRNA and a clonal barcode are encoded in the 3′ LTR such that both transcripts carry perturbation and lineage information. **B**, Library preparation and analysis strategy for PORTAL. CRISPR perturbation libraries are cloned and transduced in pooled format. Bulk mRNA is harvested at desired timepoints, and reporter and identity transcripts are amplified using a T7-mediated linear amplification strategy and quantified by sequencing. Reporter activity is normalized to identity transcript abundance within each clonal lineage, defined by a shared sgRNA and clonal barcode, yielding a distribution of phenotypes for each perturbation. **C,** Pilot screen measuring AP-1 activity following knockdown of 2,389 essential genes in K562 cells. Top panel shows the screen design using a synthetic AP-1 response element. Left panel plots the effect of each sgRNA on identity transcript expression (*x*-axis; proxy for fitness) and reporter transcript expression (*y*-axis; AP-1 activity). Because each sgRNA produces a distribution of lineage-resolved effects, effect sizes are quantified using a Mann-Whitney test relative to non-targeting control sgRNAs (orange dots; schematic top right). Red and green dots indicate sgRNAs that are outliers for AP-1 reporter activity or identity expression, respectively. Right panels show example lineage-level comparisons between selected hit sgRNAs and non-targeting controls. **D**, Validation of hit sgRNA phenotypes by flow cytometry. In the PORTAL vector, reporter activity is coupled to mCherry expression. Gating strategy shown in Fig. S1B. **E**, Gene set enrichment analysis of genes whose expression correlates with AP-1 reporter activity across perturbations in an independent Perturb-seq dataset. Enrichment was performed against transcription factor target gene sets (MSigDB C3). An AP-1 motif emerged as the top hit, confirming reporter specificity.

The nested architecture of the PORTAL vector precludes conventional PCR amplification, as recombination during amplification would generate chimeric molecules that scramble information across distinct transcripts. To preserve the linkage between reporter output, perturbation identity, and clonal barcode, we developed an amplification strategy that combines elements of Smart-seq2^56^ and CEL-seq^57^ (Methods, Fig. 1B, Fig. S1, Supplemental Protocols). During reverse transcription, a template-switching reaction introduces a T7 promoter. Following amplification by in vitro transcription, the two transcripts can be separated by size for library preparation. Reverse transcription primers also incorporate unique molecular identifiers^51^, enabling single-molecule quantification of phenotypes.

We first applied PORTAL in a pilot screen to identify modulators of AP-1 signaling. AP-1 is a dimeric transcription factor that integrates diverse upstream signals including MAPK activation, growth factors, and cellular stress^58^. Given its role as a convergent signaling node, we reasoned it would be sensitive to perturbation of many cellular processes, making it a broad test of a reporter-based screening approach. We cloned a synthetic AP-1 response element^59^ containing 7 AP-1 binding sites into our vector and used a library of 9,355 sgRNAs to knock down 2,389 essential genes^60^ (4 sgRNAs per gene when available; Supplemental Tables) in K562 cells using CRISPRi.

Sequencing detected 7 million distinct clonal lineages corresponding to unique combinations of sgRNA and clone barcode across two replicate screen arms (Fig. 1C, Supplemental Tables). To quantify perturbation effects, we regressed reporter transcript abundance against identity transcript abundance within each lineage, producing a compensated reporter signal that normalizes for lineage size, integration-site expression bias, and promoter interference (Methods, Fig. S2A). The identity transcript abundance serves as a proxy for cell number. By comparing the fold change in identity counts for each sgRNA to its abundance in the input library, we defined an inferred fitness measure that strongly correlated with the results of a previous screen (*r* = 0.64, Fig. S2B,C).

Lineage resolution allows us to measure a distribution of effects for each sgRNA, which can be compared to the distribution from cells with non-targeting control guides using a Mann-Whitney test (Methods, Fig. 1C top right, Fig. S2A,B). Replicates showed tight correlation for both compensated reporter (*r* = 0.81) and identity (*r* = 0.95) effect sizes across sgRNAs (Fig. S2D).

Plotting compensated reporter versus identity effect sizes revealed clear separation between perturbations causing general fitness defects (reduced identity expression; Fig. 1C, green), specific effects on AP-1 activity (Fig. 1C, red), and non-targeting control sgRNAs (Fig. 1C, orange). We identified perturbations that both increased and decreased AP-1 activity and validated selected hits by flow cytometry using the mCherry contained in the reporter transcript (Fig. 1D).

As orthogonal validation, we compared AP-1 reporter phenotypes to a published Perturb-seq dataset^60^ targeting the same essential genes (Methods). Gene-level reporter phenotypes were correlated with gene expression changes across perturbations to generate a ranked list, which was used as input for gene set enrichment analysis against transcription factor target gene sets. AP-1 motifs were the top enriched signatures (Fig. 1E, Fig. S2E), confirming the faithfulness of our reporter.

Having established PORTAL as a sensitive and quantitative readout, we next asked whether it could support systematic genetic interaction (GI) mapping, a substantially more demanding application that requires resolving the combined effects of perturbation pairs at scale. Meeting this challenge places new demands on both perturbation delivery and library construction.

### Mini Pol III promoters enable dual sgRNA CRISPR screens

GI mapping using PORTAL requires robust co-expression of two sgRNAs within the same cell. Past efforts to include multiple promoters within the lentiviral 3’ LTR have yielded poor performance, likely because the LTR tolerates only limited insert size before other essential functions are impaired. sgRNAs for SpCas9 are typically expressed from Pol III promoters such as human or mouse U6 (hU6 or mU6). Following past work^61^, we reasoned that Pol III promoters could be “minified” by deleting a spacer region normally occupied by a nucleosome (Fig. 2A). We designed mini promoters for five candidates—hU6, mU6, a synthetic optimized sU6^62^, an existing mini H1 construct^61^, and hU6-8 (derived from one of nine U6 copies in the human genome)—and tested their ability to drive CRISPRi knockdown of CD81 by flow cytometry. mini hU6 and mini hU6-8 performed best. We then constructed a dual-sgRNA cassette to simultaneously knock down CD81 and CD151 (Fig. 2B). Initial tests showed incomplete CD151 knockdown when the promoters both pointed downstream, suggesting promoter interference. Placing the two promoters in divergent orientation resolved this (Fig. 2B, top row), yielding strong knockdown of both targets. This cassette can be incorporated into dual-sgRNA vectors for PORTAL-based GI mapping or single-cell profiling via CROP-seq, analogous to a recent tRNA-mediated dual sgRNA approach^63^.

**Figure 2.**
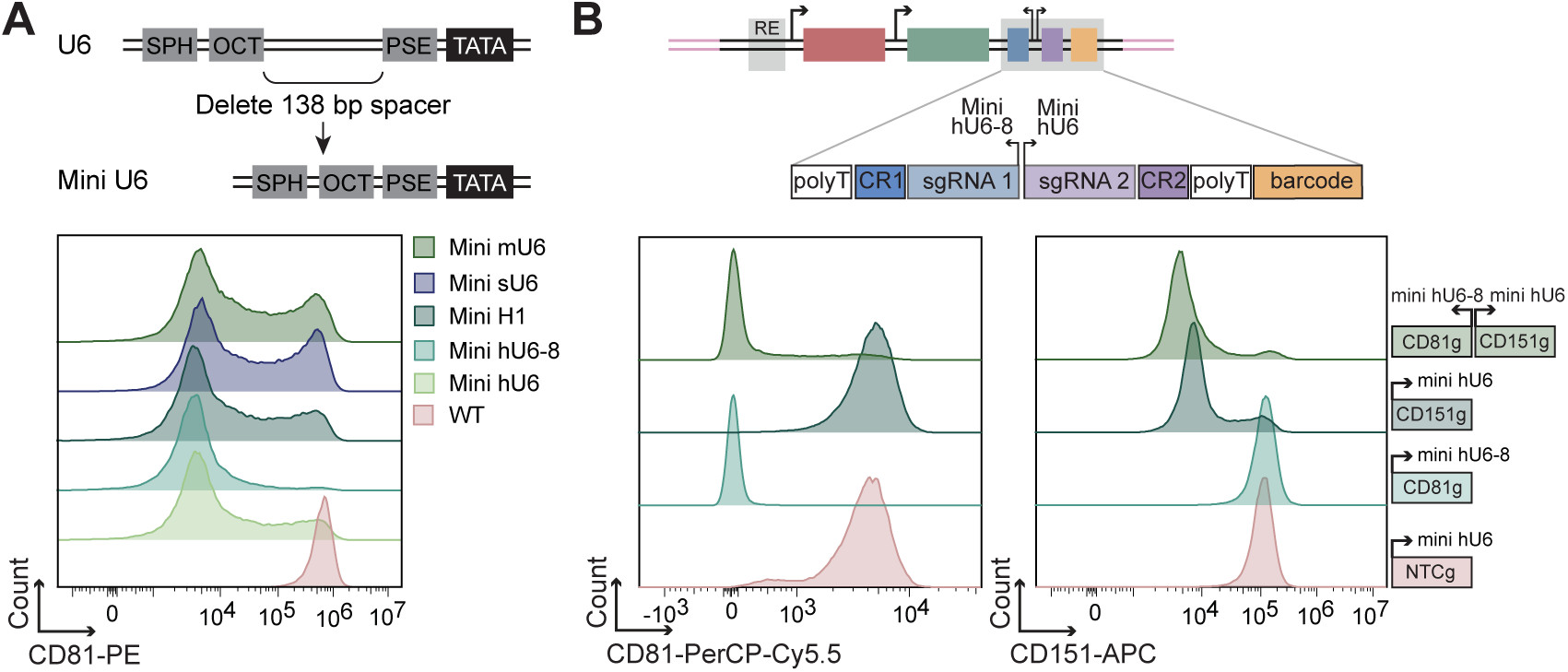
Miniaturized Pol III promoters enable efficient dual-sgRNA expression. **A**, Design of mini Pol III promoters generated by deleting a nucleosome-occupied spacer region from canonical promoters. Bottom panel shows flow cytometry quantification of knockdown of *CD81* using five candidate mini promoters. **B**, Dual-sgRNA cassette design and validation. Top row, divergent orientation of two mini promoters driving simultaneous knockdown of *CD81* and *CD151* yields robust repression of both targets. Knockdown efficiency was assessed by flow cytometry. NTC indicates a non-targeting guide.

### CAP cloning enables construction of ultrahigh diversity lentiviral libraries

The next challenge to enable PORTAL-based GI mapping is library construction. To test the limits of our approach, we designed a library targeting all pairwise combinations of 816 sgRNAs (library design described below), yielding 665,856 conditions. Combined with 120,000 sequencing error-resistant clonal barcodes^64^ (sufficient to label 1,200 lineages per sgRNA pair with a 1% collision rate) this design corresponds to approximately 80 billion possible library elements. Libraries of this diversity are difficult to clone with even representation using traditional approaches due to a bottleneck imposed by bacterial transformation.

We therefore developed CAP cloning (covalently closed assembly products), a cloning technique that bypasses bacterial transformation entirely (Methods, Supplemental Protocols). Briefly, we ordered three separate oligo libraries (sgRNA 1, sgRNA 2, and barcode), stitched these components together by PCR, and assembled the products into a lentiviral backbone. To generate sufficient material for viral packaging, the assembled library was amplified using a high-fidelity long-range polymerase. Critically, the amplification primers introduced TelN protelomerase recognition sites. Treatment with this enzyme produces covalently closed hairpin ends resistant to exonuclease digestion. These products are therefore directly competent for lentiviral packaging (Methods), consistent with prior demonstrations for individually cloned constructs^65^.

Long-read sequencing confirmed 99.8% sequence identity, comparable to conventionally cloned plasmid libraries (Fig. 3B, Fig. S3A), and revealed no increase in structural variant frequency (Fig. S3B). Deep sequencing of packaged lentiviral genomes showed good concordance between oligo synthesis representation and final library representation (Pearson *r* = 0.63; Fig. 3C), with 99.99% of all possible sgRNA pairs detected and even representation (coefficient of variation CV = 0.27; Fig. 3D). Although developed here to enable large-scale GI mapping, CAP cloning provides a general strategy for constructing ultrahigh-complexity lentiviral libraries without reliance on bacterial transformation.

**Figure 3.**
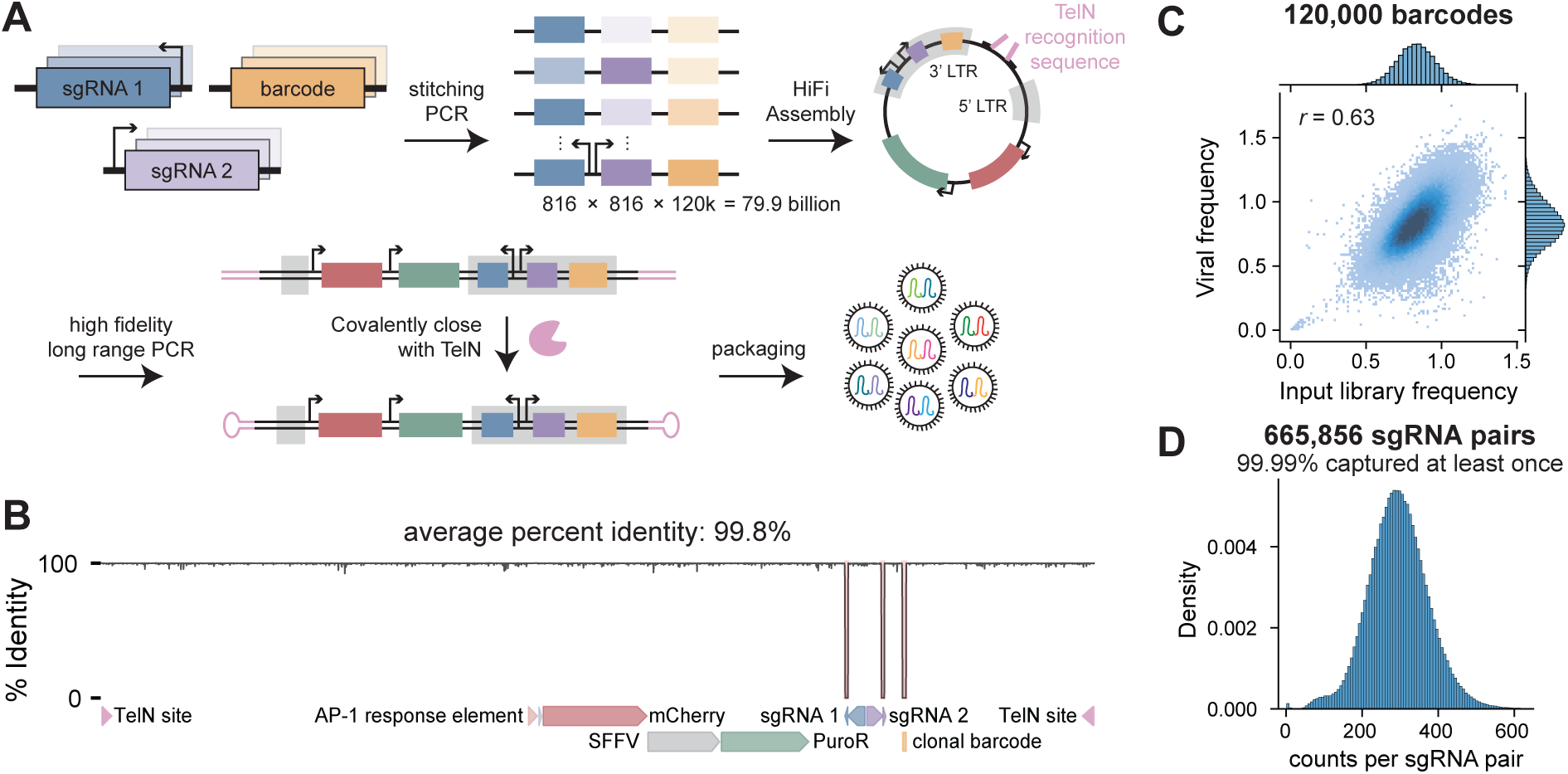
CAP cloning enables construction of ultrahigh-complexity lentiviral libraries. **A**, Overview of CAP cloning workflow used to produce a dual sgRNA library for genetic interaction mapping with PORTAL. sgRNA1, sgRNA2, and clonal barcode oligo pools were stitched together by PCR, assembled into a lentiviral backbone, and amplified with primers introducing TelN protelomerase recognition sites. TelN treatment generates covalently closed hairpin ends, producing exonuclease-resistant linear DNA molecules directly competent for lentiviral packaging without bacterial transformation. **B**, Long-read sequencing validation of CAP-cloned libraries. Plot shows percent sequence identity across the length of the PORTAL vector. Variable regions correspond to the two sgRNA protospacers and the clonal barcode. **C**, Concordance of clonal barcode representation between the oligo library used during cloning and packaged lentiviral genomes, assessed by deep sequencing. **D**, Distribution of sgRNA pair representation in packaged lentiviral genomes, showing near-complete coverage and even representation across 665,856 sgRNA pairs.

### A 665,856-condition genetic interaction map measuring AP-1 activity

We combined and applied these technologies to construct the largest exhaustive GI map measured in human cells. GI mapping aims to systematically measure how pairs of perturbations combine to influence cellular phenotypes, and can reveal functional relationships between genes, complexes, and pathways^66^. Fitness-based GI screens have been particularly successful in yeast, where exhaustive maps of pairwise interactions have provided a unifying view of cellular organization^67^. In mammalian systems, a recent study scaled GI measurements by performing genome-wide CRISPR screens across a smaller set of predefined knockout backgrounds, yielding dense interaction profiles along one axis without enumerating all pairwise combinations^34^. However, exhaustive GI screens scale quadratically with the number of target genes, and in mammalian systems fitness assays typically require 500–1000-fold cell representation per condition to resolve growth differences. Together, these constraints make exhaustive fitness-based interaction mapping challenging even at the level of cell culture in most mammalian cell types.

PORTAL offers a path around these limitations by providing direct molecular phenotypes from individual cells, substantially reducing the representation required per interaction. A key requirement, however, is that the reporter respond broadly to diverse perturbations; the appeal of fitness lies in its ability to integrate many physiological processes into a single phenotype. We hypothesized that AP-1 transcriptional activity, as a convergent node downstream of many signaling and stress pathways, would provide similarly broad sensitivity while retaining the per-cell efficiency of a reporter-based readout.

To test this hypothesis, we designed a library spanning diverse cellular processes. We mined a Perturb-seq dataset^60^ and selected 816 sgRNAs targeting 612 genes across 318 different transcriptional clusters, protein complexes, and biological processes with detectable transcriptional phenotypes (Fig. 4A, Fig. 3, Methods). Within each feature we selected at least 3 genes to provide redundancy against inactive sgRNAs.

**Figure 4.**
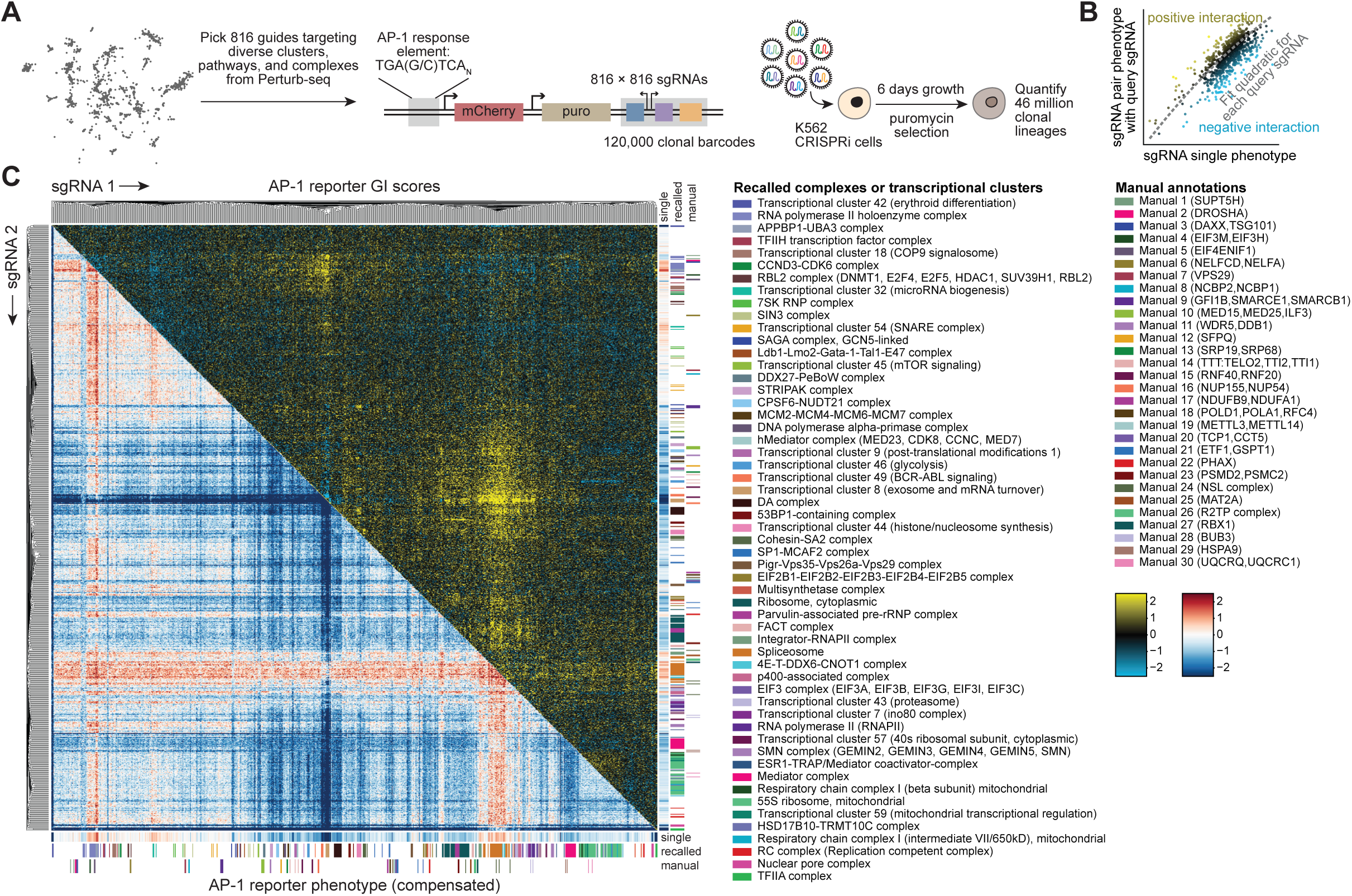
A genetic interaction map of AP-1 activity measured using PORTAL. **A**, Schematic of the genetic interaction (GI) experiment. A total of 816 sgRNAs were selected from an independent Perturb-seq dataset to target 612 genes spanning diverse transcriptional clusters, protein complexes, and biological processes. **B**, Framework for calculating GI scores. For each query sgRNA, expected AP-1 reporter activity is modeled as a quadratic function of partner sgRNA activity; deviations from this expectation define positive (synergistic) or negative (buffering) interactions. **C**, AP-1 GI map measured using PORTAL, comprising 665,856 pairwise phenotypes. The lower diagonal shows AP-1 reporter activity for each sgRNA pair, while the upper diagonal shows corresponding GI scores, which are used for clustering. The first strips along the bottom and right indicate AP-1 reporter activity for individual gene perturbations. The second strip annotates recalled biological features, including protein complexes and genes that clustered together in Perturb-seq. The third strip shows manual annotations of additional biological features or example sgRNA pairs targeting the same gene that clustered together. AP-1 reporter activity and GI scores are each scaled via the standard deviation across the dataset for visualization.

We transduced the CAP-cloned library into K562 cells and performed the screen (Methods, Supplemental Protocols). Sequencing detected 665,605 sgRNA pairs and 46 million independent clonal lineages, each providing an independent measurement of perturbation effects, with even representation across interactions (Fig. S4A, Supplemental Tables). Our cloning strategy produces each sgRNA pair in both orientations (Fig. S4B). Consistent with strong technical performance, both identity (*r* = 0.67) and AP-1 reporter (*r* = 0.68) phenotypes correlated well across orientations (Fig. S4C). This dataset is to our knowledge the largest GI map by number of targeted genes and the first at this scale using a non-fitness phenotype.

We next asked whether these data could reveal meaningful GI structure. We calculated GI scores using a framework established in prior yeast and human GI screens (Fig. 4B, Fig. S4D,E)^41,68^. For each “query” sgRNA, we defined the expected AP-1 phenotype as a function of the partner sgRNA’s phenotype by fitting a quadratic function across all pairs involving the query. Deviations from this expectation define positive (synergistic) and negative (buffering) interactions. For each sgRNA pair, we averaged measurements across both sgRNA orientations to produce final phenotype and GI score estimates.

The resulting GI map was highly structured (Fig. 4C, Supplemental Tables). To test whether this structure reflected biological function, we asked whether sgRNAs that co-clustered in the original Perturb-seq dataset (corresponding to shared transcriptional phenotypes, protein complex membership, or biological processes) also clustered in the AP-1 GI map (Methods). We observed recall of 55 features (annotations on right of Fig. 4C). The library also included 184 genes targeted by two distinct sgRNAs, of which 78 clustered together in the GI map. As in fitness-based interaction maps, these results indicate that interaction profiles derived from transcriptional reporters can be used to assign gene function.

In addition to reporter-based GIs, PORTAL also enables estimation of fitness GIs by comparing sgRNA-pair representation among observed clonal lineages to that in the input lentiviral library (Methods, Fig. S5A). Using this approach, we computed a fitness GI map from the same dataset. The fitness GI map (Fig. S5B) recalled some distinct features, demonstrating that different readouts capture different interaction types, but recalled fewer features overall. This may reflect that our screen quantified ∼70 lineages per sgRNA pair, far below typical fitness screen coverage, and supports that transcriptional reporters can extract more information per lineage.

Because our library targeted diverse cellular processes, we could examine global interaction structure. Genes within major complexes and pathways showed homogeneous interaction patterns, as expected if interaction profiles reflected shared function. This coherence allowed us to average interactions across related genes, revealing interpretable relationships between pathways. The resulting pathway-level GI map was strongly organized by biological function (Fig. S6A). For example, the tightest association was observed between perturbations targeting the DDX27-PeBoW complex, which is required for ribosome maturation, and a Perturb-seq-defined transcriptional cluster containing proteins associated with the 40S ribosome. We visualized strong pathway-level interactions on an embedding of Perturb-seq transcriptional phenotypes for the corresponding genes^60^ (Fig. S6B). For example, inhibition of glycolysis reduced AP-1 activity and strongly synergized with inhibition of multiple aspects of mitochondrial biology, including mitochondrial translation, respiratory chain function, and tRNA processing. This pattern is consistent with synthetic lethality between alternative ATP-generating pathways. Overall, these results illustrate that AP-1 activity captures functional relationships across diverse cellular processes.

Beyond global structure, GI maps can highlight specific outlier interactions that reveal functional relationships between individual genes. Perturbations inhibiting the NAE1-UBA3 complex, a core component of the neddylation pathway, showed uniformly high AP-1 reporter activity (Fig. 5A; also visible as a vertical stripe annotated as APPB1-UBA3 complex in Fig. 4C). sgRNAs targeting NEDD8, the ubiquitin-like protein attached during neddylation, were also outliers in our pilot screen (Fig. 1C). Although AP-1 activation by MLN4924, a small-molecule NAE inhibitor, has been attributed to off-target effects^69^, our genetic data support an on-target mechanism: reduced neddylation stabilizes AP-1 transcription factors, potentially through impaired turnover of c-Jun, a known substrate of Cullin RING ligases^70^ (Fig. 5A, right panel).

**Figure 5.**
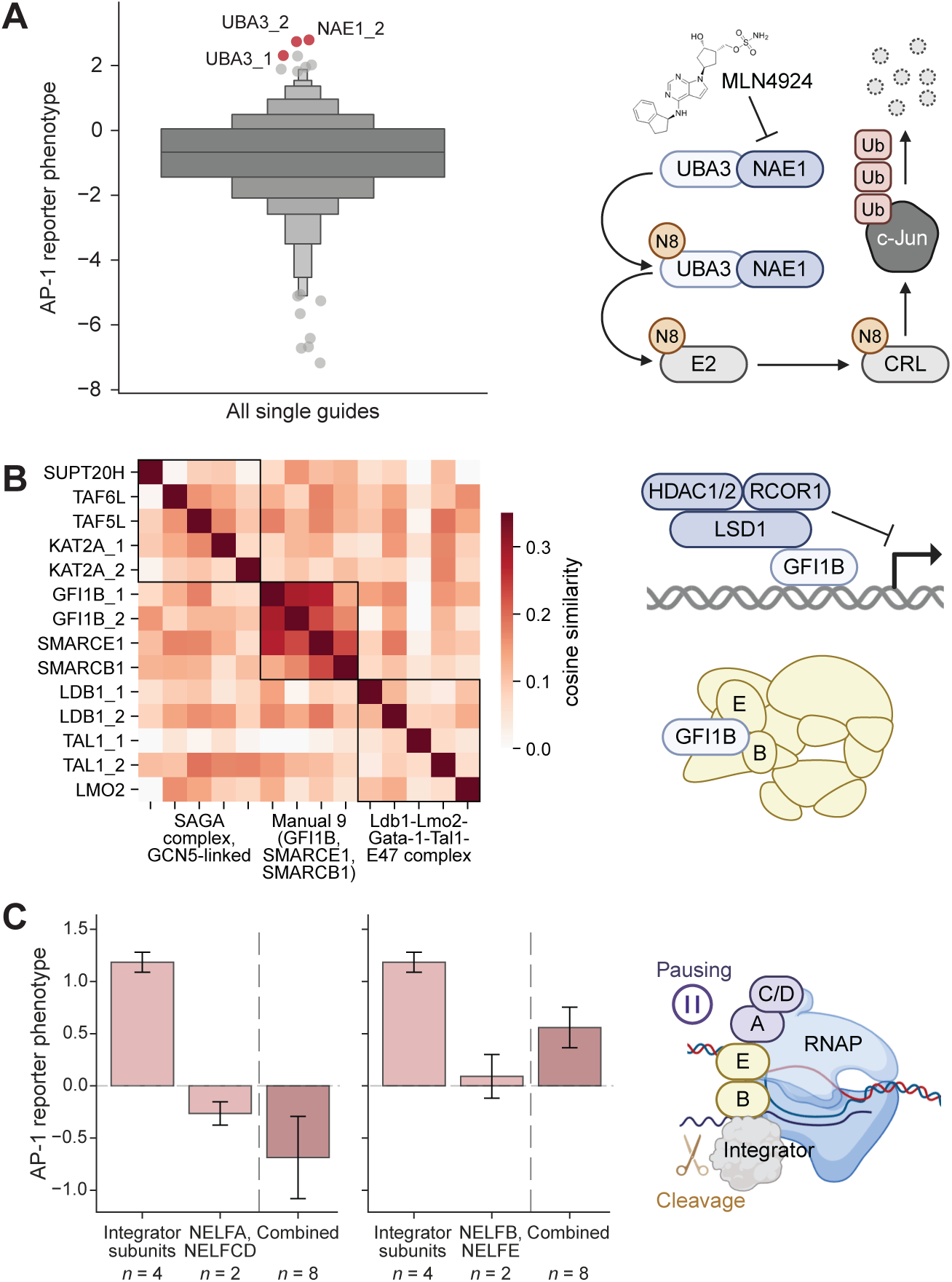
Molecular insights from the AP-1 genetic interaction map. **A**, Disruption of neddylation leads to elevated AP-1 activity. Left panel shows AP-1 reporter activity for individual sgRNAs; components of the neddylation machinery (red) are prominent outliers. Right panel illustrates a proposed mechanism in which impaired neddylation reduces Cullin-RING ligase (CRL) activity, stabilizing AP-1 transcription factors. MLN4924 is a chemical inhibitor of neddylation previously shown to activate AP-1. **B**, Clustering of GFI1B with components of the SWI/SNF chromatin remodeling complex. Left panel shows a heatmap of cosine similarity between interaction profiles for sgRNAs targeting GFI1B, SMARCE1, SMARCB1, and other sgRNAs that clustered nearby. Right panel contrasts GFI1B’s canonical LSD1-dependent repressor function with proximitome data suggesting direct interaction with SWI/SNF, potentially explaining the observed clustering. E and B denote SMARCE1 and SMARCB1, respectively. **C**, Resolution of subcomplex architecture within the NELF complex through differential interactions with Integrator subunits. Left panel shows distinct interaction patterns when Integrator knockdowns are combined with perturbations of *NELFA*/*CD* versus *NELFB*/*E*. Right panel illustrates a model in which NELF-A and -C/D promote establishment of promoter-proximal pausing, while Integrator-mediated pause release requires recruitment via NELF-B and -E. Letters indicate individual NELF subunits.

We also observed tight clustering of the transcription factor GFI1B, a key regulator of hematopoiesis, with SMARCB1 and SMARCE1, members of the SWI/SNF chromatin remodeling complex (Fig. 5B). GFI1B is best known as a transcriptional repressor that acts through its obligate effector LSD1^71^. However, proximitome studies in K562 cells have identified a physical association between GFI1B and SWI/SNF subunits that is independent of LSD1^72^. The functional clustering observed here provides orthogonal support for this interaction and suggests GFI1B and SWI/SNF may cooperate through a mechanism distinct from GFI1B’s repressor activity.

Finally, our GI map resolved subcomplex architecture within the NELF (negative elongation factor) through interactions with Integrator (Fig. 5C). These complexes act during promoter-proximal pausing, an early phase of transcription initiation: NELF helps establish the paused state while Integrator regulates its resolution by cleaving nascent transcripts^73,74^. Structural studies have shown that NELF-A and NELF-C/D form a lobe that directly engages RNA polymerase II and enforces pausing, whereas NELF-B and NELF-E form a second lobe that interacts with Integrator^75,76^. Consistent with this architecture, perturbations targeting *NELFA*/*CD* and *NELFB*/*E* exhibited distinct interaction patterns with Integrator subunits. Interactions involving NELF-A and - C/D were consistent with epistasis, in which knockdown of NELF-A and -C/D masked the effects of Integrator perturbation, consistent with the loss of paused RNA polymerase II for Integrator to release.

These examples illustrate that our AP-1 GI map captures interpretable biology at multiple scales: pathway-level organization, functional relationships between genes, and subcomplex architecture within large regulatory complexes.

### Toward comprehensive genetic interaction mapping in human cells

The technologies described above enabled measurement of 665,856 GIs. Comprehensive mapping among 10,000 genes expressed in a typical cell type would require on the order of 50 million measurements, a scale that remains challenging even with our improved efficiency. We therefore explored two strategies for further scaling.

First, single-cell resolution would further increase information yield beyond lineage- or sgRNA-level measurements. Inspired by combinatorial indexing methods^54^, we distributed cells into 96-well plates coated with oligo dT primers during RNA capture and introduced partition barcodes via uniquely barcoded reverse transcription primers in each well (Fig. 6A). The combined number of partition barcodes, clone barcodes, and sgRNA identities is sufficiently large that cells are unlikely to receive the same combination, enabling single-cell resolution (Methods). As a proof of concept, we profiled interactions among 100 hit genes from our pilot experiment (Fig. 6A, Supplemental Tables). Sequencing identified ∼20 million distinct cells across ∼4.6 million clonal lineages and 10,000 sgRNA pairs. Fig. 6B shows example distributions of reporter expression at the single-cell level within clonal lineages for sgRNA pairs that induced (simultaneous knockdown of *KDM5C* and *NEDD8*) or repressed (*MTOR* and *RNF20*) AP-1 activity, spanning an approximately 30-fold dynamic range. Beyond increasing statistical power, single-cell resolution enables new applications, such as screens for perturbations that regulate cell-to-cell variability, which may be particularly relevant for understanding cell fate decisions or incompletely penetrant phenotypes in drug resistance.

**Figure 6.**
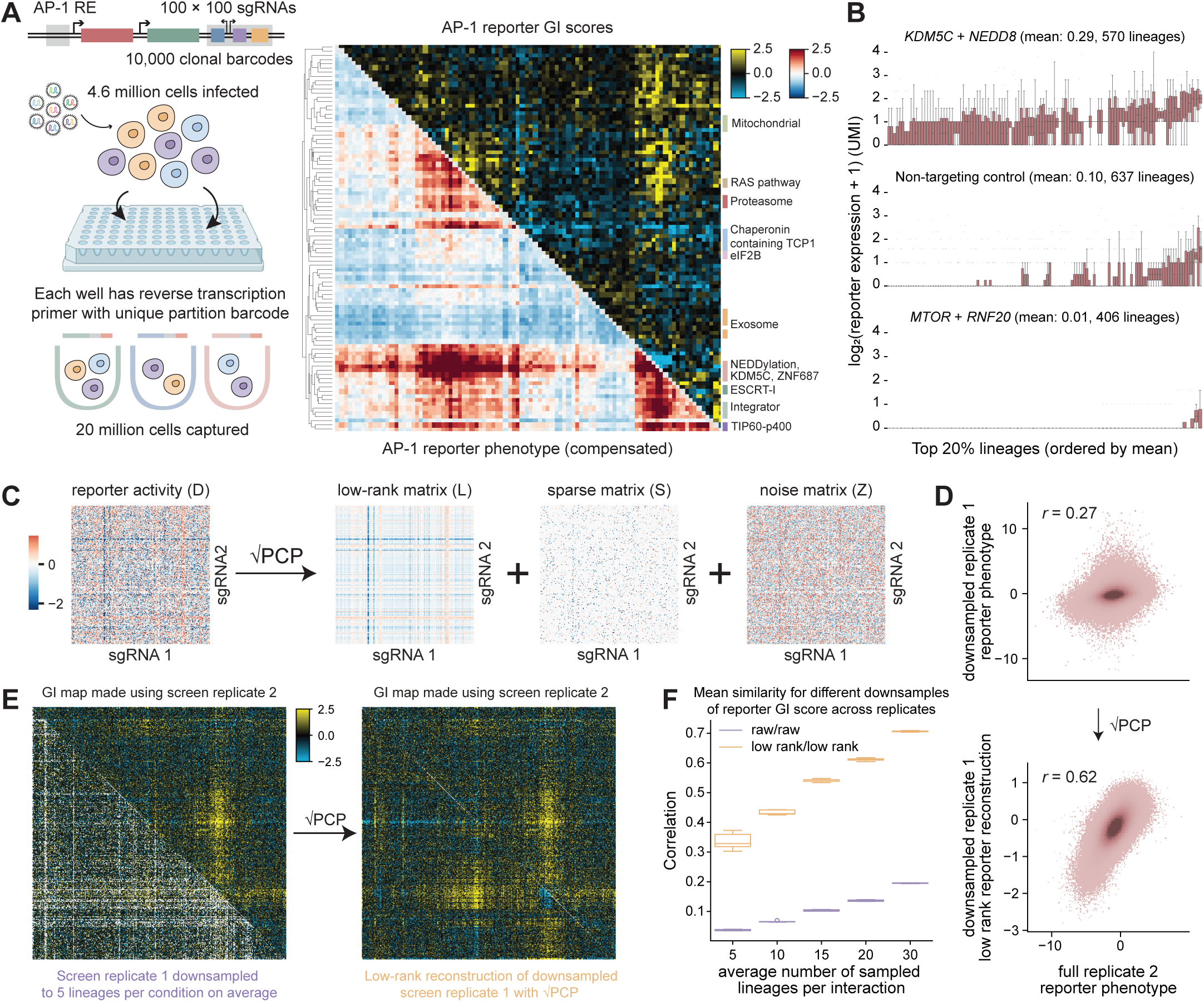
Paths toward comprehensive genetic interaction mapping. **A**, Schematic of single-cell PORTAL using combinatorial indexing. Cells are distributed across wells and reverse transcription is performed with barcoded primers, enabling single-cell resolution when combined with sgRNA and clonal barcode diversity. Right panel shows AP-1 reporter activity (lower diagonal) and GI scores (upper diagonal) measured at single-cell resolution for interactions among 100 hit sgRNAs from the pilot screen. B, Example single-cell distributions of AP-1 reporter activity within clonal lineages. Box plots show AP-1 reporter activity for cells within the top 20% of clonal lineages (ranked by mean activity) for three example sgRNA pairs exhibiting high, unperturbed, or low reporter activity. **C**, Schematic of square root principal component pursuit (√PCP) for matrix completion, which decomposes observed measurements into a low-rank component capturing large-scale structure, sparse outliers, and noise. **D**, Recovery of structure in AP-1 reporter activity from sparsely sampled measurements. Lineages from the GI map in Fig. 4 were split into two replicates, and the first replicate was downsampled to an average of five lineages per sgRNA pair. AP-1 reporter activity for sgRNA pairs was compared between replicates before (top) and after (bottom) √PCP reconstruction. Each point represents an sgRNA pair. **E**, GI maps computed from raw measurements and from √PCP reconstructions mirror trends from (d). Left panel compares GI maps derived from downsampled replicate 1 (lower diagonal) and fully sampled replicate 2 (upper diagonal). Right panel shows that √PCP reconstruction recovers large-scale interaction structure. **F**, Low-rank reconstruction improves cross-replicate reproducibility of GI maps. Each replicate was independently downsampled 5 times to different average numbers of lineages per sgRNA pair and used to compute GI maps. Correlation between replicate GI maps is plotted as a function of sampling depth for raw measurements (purple) and √PCP-reconstructed maps (orange).

Second, computational imputation may enable reconstruction of GI structure from sparse measurements^77^. Empirically, GI matrices exhibit low-rank structure, meaning large-scale patterns such as complex- or pathway-level interactions can be captured by a smaller number of underlying components. We therefore explored square root principal component pursuit (√PCP)^78^, a matrix completion method that separates observations into a low-rank matrix capturing large-scale structure, a sparse matrix capturing outliers, and noise (Fig. 6C). Because √PCP is hyperparameter-free, it is well-suited to settings where ground truth is unknown.

To test whether √PCP can recover GI structure under limited sampling, we randomly split the lineages in our GI map (Fig. 4C) into two replicates. We downsampled the first replicate to an average of five lineages per sgRNA pair and compared compensated reporter expression to the second replicate, which retained an average of 35 lineages per pair, yielding modest agreement (*r* = 0.27; Fig. 6D, top). Applying √PCP to the downsampled replicate substantially improved correspondence with the full replicate (*r* = 0.62; Fig. 6D, bottom), and the resulting GI maps exhibited visually similar large-scale structure (Fig. 6E). As an internal validation, we independently downsampled and reconstructed both replicates. √PCP markedly increased cross-replicate reproducibility (Fig. 6F), with agreement improving with number of lineages, indicating convergence toward shared low-rank structure. Together these results suggest that low-rank GI structure can be reconstructed from sparsely sampled measurements, pointing toward a combined computational-experimental path to genome-scale GI mapping.

## Discussion

We have introduced two technologies—PORTAL and CAP cloning—that together enable pooled CRISPR screens at scales and sensitivities beyond what existing methods can achieve. In its simplest configuration, PORTAL can be used without a reporter to perform lineage-resolved CRISPR fitness screens, which is particularly valuable in positive-selection contexts where clonal jackpotting is a major confounder. When coupled to transcriptional reporters, PORTAL captures quantitative phenotypes with single-molecule resolution that extend beyond fitness effects. In this study, AP-1 emerged as a broad integrator of perturbation effects, capturing interpretable biology across diverse cellular processes. Other pathways with convergent signaling properties may prove similarly useful. Looking forward, we anticipate that computational design of synthetic regulatory elements will further expand the space of viable reporter phenotypes^79,80^. Indeed, by jointly varying large libraries of reporter promoter sequences and genetic perturbations within a single experiment, future implementations of PORTAL could themselves function as massively parallel assays for learning sequence-to-activity relationships, generating the data needed to constrain models of promoter function and design effective reporters *de novo*.

Although we developed CAP cloning to enable large-scale GI mapping, its utility extends well beyond. Any application requiring lentiviral delivery of highly diverse libraries stands to benefit, including massively parallel reporter assays, large-scale lineage tracing experiments, pooled screening of computationally designed sequences, constructing ORF libraries encoding proteins toxic to bacteria, combinatorial assembly of proteins such as antibody variable region pairings, and high-complexity single-cell screens to provide training data for emerging virtual cell models.

Finally, our results outline a path toward comprehensive GI mapping among all human genes. The screen reported here—665,856 pairwise perturbations measured across 46 million clonal lineages—demonstrates that large-scale GI mapping with transcriptional reporters is feasible.

Extending this approach to interactions among all expressed genes in a cell type would require measuring on the order of 2.5 billion lineages, an increase of less than two orders of magnitude. In experimentally tractable suspension cell lines, brute-force strategies may be sufficient, as the required cell culture and scale of sequencing lies within existing infrastructure. In systems where such scaling is impractical, computational reconstruction from sparse measurements offers a complementary route to recover pathway- and complex-level relationships. Whether through experimental scale, computational assistance, or a combination of both, comprehensive maps of human GIs appear within reach.

## Supporting information

Supplemental Protocols

## Acknowledgments

We gratefully acknowledge all the members of the Norman Lab for their valuable thoughts and discussion on this work. We acknowledge the use of the Integrated Genomics Operation Core and the Flow Cytometry Core Facility, which are funded by the NCI Cancer Center Support Grant (CCSG, P30 CA08748), Cycle for Survival, and the Marie-Josée and Henry R. Kravis Center for Molecular Oncology. This work was funded by NIH R21 HG012230 and NIH Director’s New Innovator Award DP2 GM140925 (to T.M.N.). R.C.A. is a Robert Black Fellow of the Damon Runyon Cancer Research Foundation (DRG-2462-22).

## Author contributions

Conceptualization, A.T., R.C.A., and T.M.N. Methodology, A.T., R.C.A., R.E.M., and T.M.N. Formal analysis, A.T. Validation, R.C.A and R.E.M. Investigation, A.T., R.C.A., and T.M.N. Writing – Original Draft, A.T., R.C.A., and T.M.N. Visualization, A.T., R.C.A., and T.M.N. Supervision, T.M.N. Funding Acquisition, T.M.N.

## Data Availability

Raw sequencing data and metadata are available on the Sequence Read Archive with accession [to be deposited]. Processed guide-level phenotypes for PORTAL screens are in Supplemental Tables. Lineage-level or single-cell count data for PORTAL screens are available on Zenodo: [to be deposited] (pilot screen targeting essential genes, Fig. 1), [to be deposited] (genetic interaction map, Fig. 4), and [to be deposited] (single-cell PORTAL screen, Fig. 6). External K562 Perturb-seq datasets: https://gwps.wi.mit.edu/.

## Code Availability

All custom analysis code generated in this study is available at https://github.com/ https://github.com/norman-lab-msk/PORTAL.

## Experimental Methods

### Cell culture

K562 (ATCC, CCL-243) cells were purchased from American Type Culture Collection (ATCC). Lenti-X™ 293T Cell Line was purchased from Takara Bio (632180). K562 KRAB-dCas9 is a gift from Jonathan Weissmann. K562 were maintained in RPMI-1640 (Gibco, 11875135) supplemented with 10% FBS and Pen/Strep. Lenti-X™ 293T is maintained in DMEM (Gibco, 11965092) with 10% FBS and Pen/Strep. Cells are passaged with CTS™ TrypLE™ Select Enzyme (Thermo Fisher, A1285901). Cell counting was done using the Countess® II FL Automated Cell Counter.

### Plasmid designs

pRCA381 - pHR-UCOE-SFFV-ZIM3-dCas9-P2A-mCherry was cloned from pJB108 by replacing the BFP with mCherry. pBA950 was used as the backbone to clone pScreen backbone. pAT005 was cloned from pBA950 by removing WPRE, replacing BFP with GFP, inserting AP1-mCherry cassette, replacing 3’ LTR with 3’ LTR single guide, and replacing EF-1a with SFFV. 3’ LTR single guide consists of a mU6 promoter, a lacZ fragment, sgRNA constant region, a barcode region, and a Nextera Read 2 sequence in reverse orientation for direct RT from the transcripts. pAT011 was cloned from pAT005 by removing the mU6 promoter and replacing it with sgRNA constant region 1 in reverse orientation. pRCA944 was cloned from pAT011 by removing GFP and replacing it with PuroR with BsmBI site silently mutated. DNA sequences are in Supplemental Tables (response element, mCherry, GFP, 3’ LTR single guide, PuroR).

### Cloning of single-guide library

pAT005 backbone vector was digested with FastDigest BstXI (Thermo Scientific, FD1024) and BamHI (Thermo Scientific, FD0054) and purified using 0.88X ProNex Size-Selective Beads (Promega, NG2001).

The essential genes guide library and 10k barcode library were ordered from Twist Biosciences. Sequences for both libraries are in Supplemental Tables. The libraries were amplified using 8 cycles of KAPA HiFi HotStart ReadyMix PCR (Roche, 07958935001). To prepare them for stitching, primers were designed with overhangs to introduce the sgRNA constant region as overlapping ends. 5 cycles of stitching PCR were performed to join the overlapping products. Primers that bind to the start and end of the full-length product were added and 5 final cycles of PCR were performed to amplify the correctly stitched full-length product. The product was purified using 1X-2X(+1X) ProNex Size-Selective Beads (Promega, NG2001).

The stitched full-length product was ligated into the digested backbone using NEBuilder® HiFi DNA Assembly Master Mix (NEB, E2621). To remove any undigested backbone, Bsu36I (NEB, R0524S) restriction enzyme was added, incubated at 37 °C for 15 minutes and heat inactivated at 80 °C for 20 minutes. The reaction was purified using Monarch® PCR & DNA Cleanup Kit (NEB, T1030).

35 cycles of long-range PCR were performed on the ligation product using repliQa HiFi ToughMix® (Quantabio, 95200). Primers were designed to amplify the entire lentiviral genome with overhangs to introduce TelN recognition sites. The product was gel purified using QIAquick Gel Extraction Kit (Qiagen, 28704). Purified product was treated with TelN Protelomerase (NEB, M0651) and purified using Monarch® PCR & DNA Cleanup Kit (NEB, T1030).

### Cloning of dual-guide libraries

pRCA944 or pAT011 backbone vector was digested with Esp3I (NEB, R0734S) and Bsu36I (NEB, R0524S) and purified using 0.88X ProNex Size-Selective Beads (Promega, NG2001).

The sgRNA1 guide library, sgRNA2 guide library, and 120k barcode library were ordered from Twist Biosciences. Sequences for libraries are in Supplemental Tables. The libraries were amplified 8 cycles using KAPA HiFi HotStart ReadyMix PCR (Roche, 07958935001). To prepare them for stitching, primers were designed with overhangs to introduce the mini promoters and sgRNA constant region 2 as overlapping ends. 5 cycles of stitching PCR were performed to join the overlapping products. Primers that bind to the start and end of the full-length product were added and 5 final cycles of PCR were performed to amplify the correctly stitched full-length product. The product was purified using 1.25X ProNex Size-Selective Beads (Promega, NG2001).

The stitched full-length product was ligated into the digested backbone using NEBuilder® HiFi DNA Assembly Master Mix (NEB, E2621). The reaction was purified using Monarch® PCR & DNA Cleanup Kit (NEB, T1030).

35 cycles of long-range PCR were performed on the ligation product using repliQa HiFi ToughMix® (Quantabio, 95200). Primers were designed to amplify the entire lentiviral genome with overhangs to introduce TelN recognition sites. The product was gel purified using QIAquick Gel Extraction Kit (Qiagen, 28704). Purified product was treated with TelN Protelomerase (NEB, M0651) and purified using Monarch® PCR & DNA Cleanup Kit (NEB, T1030).

### PacBio long read sequencing

To produce a plasmid library for comparison, HiFi-assembled product from the dual-guide library cloning was electroporated with an Eppendorf Eporator into Endura Electrocompetent Cells (LGC, 60242-2) according to manufacturer’s protocol. Overnight cultures are grown at 32°C for 18 hours before midiprep. Midiprep was done using the QIAGEN Plasmid Plus Midi kit (QIAGEN, 12945).

Plasmid library was digested with SspI (NEB, R3132) and the cut product was size selected for 7500bp using the BluePippin 0.75% DF Low Voltage 1-6kb Marker S1 (SAGE SCIENCE, BLF7510). For CAP cloning, RepliQA amplified product from dual guide CAP cloning step 5 (Supplemental Protocols) was size selected as above targeting 7000bp. The two products were sent to Azenta for PacBio long-read amplicon sequencing on the Revio system.

### Lentiviral packaging

Lentivirus was produced by co-transfecting LentiX™ 293T cells with transfer plasmids and standard packaging vectors (psPAX2, pMD2G) using TransIT-LTI Transfection Reagent (Mirus, MIR 2300). When required, virus supernatant was concentrated using the Lenti-X™ concentrator (Takara Bio, 631231).

### Lentiviral packaging of CAP cloned product

Lentivirus was produced by co-transfecting LentiX™ 293T cells with 400ng of CAP cloned product and standard packaging vectors (366ng psPAX2, 183ng pMD2G) using TransIT-LTI Transfection Reagent (Mirus, MIR 2300) per 6-well well.

### Cell line generation

To generate new CRISPRi K562 cells (hereon named K562 ZIM3-dCas9), 1×10^6^ WT K562 was transduced with 500µL of a 5x concentrated pRCA381 viral supernatant with 8ug/mL Polybrene (Millipore Sigma, TR-1003). In general, spinfection for K562 was done at 800g for 45 minutes. One day post spinfection, cell culture media is exchanged for growth media. We generated single cell clones of K562 ZIM3-dCas9 by sorting single cells into 96-well plates and waited for outgrowth.

### Vector knockdown validation by flow cytometry

For testing guide vectors, 1×10^6^ K562 KRAB-dCas9 cells were transduced with guide vector virus aiming for a ∼10% transduction efficiency. At day 2 post transduction, GFP+ cells were sorted for purity. Day 7 post transduction, cells were spun down, washed and stained with stained on ice for 30 minutes in the dark. Flow cytometry was performed using a BD LSRFortessa or Thermo Fisher Attune cytometer, with data analysis conducted in FlowJo. Parental lines expressing GFP, BFP, or mCherry were used as single-color compensation controls. Antibodies used included CD81-PE (Biolegend, 349505 1:25), CD81-PerCP-Cy5.5 (BD, 565430 1:25), and CD151-PE (BD 556057, 1:10)

### Single guide pilot screen validation

1×10^6^ K562 KRAB-dCas9 cells were spinfected with lentivirus of pAT005 with guides targeting KDM5C, GAB2, or non-targeting guide. One day post spinfection, media was exchanged for fresh growth media. Cells were passaged every two days in growth media. On day 6 post transduction, cells were analyzed on an Thermo Fisher Attune cytometer.

### Viral supernatant library to assess genetic interaction map library representation

RNA from viral supernatant of packaged CAP-cloned genetic interaction library was collected using the Quick-RNA Viral Kit (Zymo R1034) per manufacturer’s protocol. Reverse transcription was performed using the Template Switching RT Enzyme mix (NEB M0466) and indexed RT primers (pScreen_7xx; sequences in Supplemental Tables). RT reaction conditions were 42°C for 90 minutes, 85°C for 5 minutes, and 4°C until further processing. A 3X ProNex size selection was performed to exchange buffers. Sequencing adapter PCR was performed using NEBNext® Ultra™ II Q5® Master Mix with P7 and pScreen_dual_5xx primers, followed by two-sided purification with ProNex beads at a 1.0x-1.3x ratio. Sequencing library was eluted in 22uL TE buffer. Quantification was performed using Qubit™ 1X dsDNA High Sensitivity (Invitrogen, Q33231), and quality control by Agilent BioAnalyzer High Sensitivity DNA Kit (5067-4626). The library was sequenced on an AVITI2 × 150 bp Cloudbreak FS Low Output flow cell (Element Biosciences, 860-00011), with a read structure of I1:16, I2:8, R1: 60, R2: 172, yielding ∼250 million reads. Bases2Fastq (Element Biosciences) was used to generate FASTQ files.

### Twist 120k barcode representation library (for comparison to previous)

Input library was prepared by 6 cycles of PCR amplification using NEBNext® Ultra™ II Q5® Master Mix (NEB, M0544S) from the Twist 120k barcode library with primers containing Illumina P5 and P7 adapters and Nextera i5, i7 indices. Library was purified using 2X ProNex Size-Selective Beads (Promega, NG2001). Following quantification using Qubit™ 1X dsDNA High Sensitivity (Invitrogen, Q33231) and quality control by Agilent BioAnalyzer High Sensitivity DNA Kit (5067-4626), the library was sequenced on an AVITI 2 × 75 bp Cloudbreak FS Medium Output flow cell (Element Biosciences, 860-00014), with a read structure of I1:16, I2:8, R1: 113, R2: 28, yielding ∼140 million reads. Bases2Fastq (Element Biosciences) was used to generate FASTQ files.

### CAP cloning library representation (pilot screen)

The validation library for the single-guide pilot screen was prepared by PCR amplification using NEBNext® Ultra™ II Q5® Master Mix (NEB, M0544S) from the CAP cloning product with primers containing Illumina P5 and P7 adapters and Nextera i5, i7 indices. Library was purified using 2X ProNex Size-Selective Beads (Promega, NG2001). Following quantification using Qubit™ 1X dsDNA High Sensitivity (Invitrogen, Q33231) and quality control by Agilent BioAnalyzer High Sensitivity DNA Kit (5067-4626), the library was sequenced on MiSeq yielding ∼4 million 50 bp paired-end reads.

### Single-guide pilot screen (Figure 1)

K562 KRAB-dCas9 cells were transduced with pAT005-essentials lentiviral library at a multiplicity of infection of ∼15%. 30 million cells were transduced across two replicates using spinfection at 800 x g for 2 hours at 30°C in the presence of 8 μg/mL Polybrene (Millipore Sigma, TR-1003). Three days post-transduction, GFP-positive cells were isolated by fluorescence-activated cell sorting. Sorted cells were cultured for an additional 3 days and harvested on day 6 post-transduction at approximately 40 million cells per replicate. Cell pellets were frozen at −80°C and subsequently processed for bulk library preparation.

### Genetic interaction map (Figure 4)

K562 ZIM3-dCas9 cells were transduced with 10X concentrated pAT011-dual816 lentivirus (targeting a 0.2-0.3 MOI). 225 million cells spread across 12 6-well plates (3 million cells per well) were spinfected with 100uL 10X concentrated virus per well at 800g for 45 minutes with 8 μg/mL Polybrene in triplicates (Total 675 millioncells). One day post transduction, each replicate was spun down and resuspended into 1 T175 flasks with fresh media. Day 2 post transduction, cells were split into 2 T175 and growth media was replaced with 1.5ug/mL Puromycin media. Day 4 post transduction, cells were spun down to remove dead cells and debris and puromycin media was refreshed. On day 5, cells were spun down with each flask were split into two T175s replacing puromycin media with growth media for recovery. On day 6 post transduction, cells were harvested and prepared for bulk library preparation.

### Single-cell genetic interaction map (Figure 6)

K562 KRAB-dCas9 cells were transduced with pAT011-dual100 lentiviral library at a multiplicity of infection of ∼7.5%. 72 million cells were transduced across two replicates using spinfection at 800 x g for 2 hours at 30°C in the presence of 8 μg/mL Polybrene (Millipore Sigma, TR-1003). Three days post-transduction, GFP-positive cells were isolated by fluorescence-activated cell sorting. Sorted cells were cultured for an additional 3 days and harvested on day 6 post-transduction. Cells were directly processed for mRNA isolation and library preparation using the single-cell library preparation protocol.

### Bulk library preparation

Total RNA was extracted from cell pellets using QIAGEN RNeasy® Midi kit (Qiagen, 75144). mRNA was isolated using PolyATtract® mRNA Isolation Systems III (Promega, Z5300).

Template-switching first strand cDNA was synthesized using Maxima H Minus Double-Stranded cDNA Synthesis Kit (Thermo Scientific, K2562). RT primers (pScreen_7xx) bind to the Nextera Read 2 sequence downstream of barcode and introduces Nextera i7 indices and UMI. In addition, a template-switching oligo TN006-TSO_T7 was used to introduce a T7 promoter. First strand synthesis was performed at 42°C for 90 min.

T7 RNA synthesis was performed directly from first strand cDNA using HiScribe® T7 Quick High Yield RNA Synthesis Kit (NEB, E2050L) at 42°C for 2 hours. DNAseI was added and incubated at 37°C for 15 minutes. RNA was purified using 1.8X RNAClean XP (Beckman Coulter, NC0068576).

A second RT reaction was performed with P7 primer at 50°C for 45 min. First strand cDNA was purified using 3X ProNex Size-Selective Beads (Promega, NG2001). Second strand synthesis was performed at 16°C for 2 hours. Reactions were treated with RNAseI to remove residual RNA. Double-stranded cDNA was purified using ProNex beads.

Identity and reporter transcripts were separated using BluePippin 0.75% DF Low Voltage 1-6kb Marker S1 (SAGE SCIENCE, BLF7510). Reporter transcript was pre-amplified 8 cycles using repliQa HiFi ToughMix® (Quantabio, 95200) with a primer that binds in the mCherry sequence and P7. Identity transcript proceeded directly to final library amplification.

Final PCR amplification was performed using NEBNext® Ultra™ II Q5® Master Mix (NEB, M0544S) with Nextera i5 indexed forward primers and P7 for 8 cycles (identity transcript) or 6 cycles (reporter transcript). Products were purified and size-selected using ProNex beads. If product amount was insufficient for sequencing, additional P5/P7 amplification was performed using NEBNext® Ultra™ II Q5® Master Mix for 4 cycles, followed by purification with ProNex beads. Quantification was performed using Qubit™ 1X dsDNA High Sensitivity (Invitrogen, Q33231), and quality control by Agilent BioAnalyzer High Sensitivity DNA Kit (5067-4626).

### Single-cell library preparation

mRNA was isolated from 250,000 cells in 30 µL per well using mRNA Catcher™ PLUS 96-well plates (Invitrogen, K157002) according to manufacturer’s protocol. Two replicates were prepared with 24 wells each.

Template-switching first strand cDNA was synthesized using Maxima H Minus Double-Stranded cDNA Synthesis Kit (Thermo Scientific, K2562). Well-specific RT primers that bind to the Nextera Read 2 sequence downstream of barcode with indexing and UMI, and template-switching oligo TN006-TSO_T7 were added directly to wells. First strand synthesis was performed at 42°C for 90 min.

cDNA was pooled by column and T7 RNA synthesis was performed directly from first strand cDNA using HiScribe® T7 Quick High Yield RNA Synthesis Kit (NEB, E2050L) at 42°C for 2 hours. DNAseI was added and incubated at 37°C for 15 minutes. RNA was purified using 1.8X RNAClean XP (Beckman Coulter, NC0068576). Subsequent steps followed the bulk library prep protocol from second RT reaction through final PCR amplification.

### Sequencing of screen libraries

#### Single-guide pilot screen

The libraries were prepared through the bulk library preparation protocol and sent to the MSKCC Integrated Genomics Operation core facility for sequencing. After PicoGreen quantification and quality control by Agilent BioAnalyzer, libraries were pooled equimolar and run on a NovaSeq 6000 in a PE50 run, including 16 cycles for the Index1 read and 8 cycles for the Index2, using the NovaSeq 6000 S2 Reagent Kit (100 Cycles) (Illumina). The loading concentration was 0.7 nM and a custom sequencing primer (seq_primer) was used instead of the standard Illumina Sequencing Primer Mix for Read1. As a result of the custom primer, the PhiX control library was not used. The run yielded ∼2.1 billion reads.

#### Dual-guide genetic interaction map

The libraries were prepared through the bulk library preparation protocol and pooled equimolar. Four sequencing runs were performed on 2 × 150 bp Cloudbreak FS High Output flow cells (Element Biosciences, 860-00013), with a read structure of I1:16, I2:8, R1: 60, R2: 172, yielding a total of ∼4 billion reads. Bases2Fastq (Element Biosciences) was used to generate FASTQ files.

#### Single-cell genetic interaction map

The libraries were prepared through the single-cell library preparation protocol and pooled equimolar. The library was then processed through the Element Adept Library Compatibility Workflow using the Adept Rapid PCR-Plus Kit (Element Biosciences, 830-00018) and sequenced on an AVITI 2 × 150 bp Cloudbreak High Output flow cell (860-00003), with a read structure of I1:16, I2:8, R1: 60, R2: 172, yielding ∼500 million reads. Bases2Fastq (Element Biosciences) was used to generate demultiplexed FASTQ files.

### Computational Methods

#### Clonal barcode library design

Pre-generated error-correcting DNA barcodes were downloaded from the freebarcodes^64^ GitHub repository. Two barcode files were concatenated to create 25bp barcodes: a 15bp barcode set correcting up to 2 errors (barcodes15-2.txt) and a 10bp barcode set correcting up to 1 error (barcodes10-1.txt). Concatenated barcodes were filtered to remove those with hairpin melting temperatures exceeding 20°C (calculated using primer3), restriction enzyme recognition sites for BstXI, BlpI, BamHI, and SalI, or the polyA signal sequence AATAAA. After filtering, barcodes were randomly ordered and the first 10,000 were selected for the single-guide pilot and single-cell screen libraries, and the first 120,000 for the dual-guide screen library. For library synthesis by Twist Bioscience, left and right adapter sequences were added to enable PCR amplification and HiFi ligation.

#### Design of essentials targeting guide library for pilot screen

We targeted 2545 genes identified as essential in a previous Perturb-seq dataset knocking down essential genes in RPE-1 cells^60^. This experiment used one dual guide construct per target, yielding the first two guides in the library. We added two additional guides per target using CRISPick. 100 non-targeting control guides also taken from the Perturb-seq experiment were added at 4X representation.

#### Design of genetic interaction map library

We designed a focused guide library targeting 612 genes to enable construction of a genetic interaction map using our AP-1 reporter. Gene selection was guided by two objectives: (1) representing diverse biological processes to demonstrate broad reporter responsiveness, and (2) including multiple perturbations per functional feature to enable validation of the method by clustering of co-functional genes.

Candidate genes were drawn from a genome-wide Perturb-seq experiment in K562 cells, which used dual-guide constructs. We restricted selection to genes with evidence of perturbation activity based on the number of differentially expressed genes detected (>50 genes at Benjamini-Hochberg corrected significance), sufficient cell recovery (>25 cells), and evidence of target knockdown or fitness effects.

Genes were selected through complementary strategies: (1) manually curated exemplars from 64 transcriptional phenotype clusters representing different biological processes; (2) systematic sampling from CORUM protein complexes with ≥75% subunit coverage in our targetable set; (3) a greedy set-cover algorithm to maximize complexes with complete representation; and (4) supplemental sampling of large complexes (≥9 subunits) to ensure ≥33% coverage of structures such as the mitochondrial ribosome, respiratory chain complex I, and spliceosome.

For guide selection, we chose the guide in the first position of the original dual-guide constructs; because of this architecture we could not definitively attribute phenotypes to individual guides. For 184 genes—those appearing in ≥7 well-represented complexes or members of small (≤3 member) perturbed complexes—we included two independent guides to provide internal replication. The final library comprised 796 targeting guides plus 20 non-targeting control guides selected for minimal transcriptional and fitness effects in prior screens.

#### Design of single-cell genetic interaction map library

Guides for the single-cell genetic interaction screen were selected from statistically significant reporter phenotype hits in the pilot screen. We selected guides showing strong reporter phenotypes (either increased or decreased) that were not among the most extremely depleted for the identity phenotype. Non-targeting control guides were selected from controls showing no significant reporter or identity phenotype in the pilot screen. For each gene, we selected the single guide with the strongest statistical significance. The final libraries contained 93 reporter hits and 7 non-targeting control guides for pairwise combinations.

#### Cloning product fidelity assessment by PacBio sequencing

##### Read alignment and processing

Protospacer and barcode regions in the reference sequence were masked prior to alignment. PacBio HiFi reads in FASTQ format were aligned to the masked reference using minimap2 v2.30-r1287 with the map-hifi preset. Alignments were converted to BAM format, sorted, and indexed using samtools v1.21.

##### Structural variant detection

To identify reads containing structural variants (SVs), we used Sniffles2 v2.6.3 with highly sensitive parameters to capture low-frequency variants. Variants were detected with minimum length of 10 bp and minimum support of 1 read, with no mapping quality filter applied. Mosaic variant detection was enabled to identify variants present in mixed read populations.

##### Per-base identity calculation

To ensure accurate per-base identity measurements, reads supporting structural variants were removed from the analysis. Reads were extracted from the VCF RNAMES field and filtered from the alignment to generate a BAM containing only reads without structural abnormalities. Per-base sequence identity was calculated from the filtered BAM files by parsing CIGAR strings and MD tags to determine matches, mismatches, insertions, and deletions at each reference position. Identity was computed as the percentage of aligned bases matching the reference sequence. Protospacer and barcode regions were excluded from the analysis.

#### Viral supernatant library to assess genetic interaction map library representation

##### Input library barcode quantification

FASTQ files from paired-end sequencing of the synthesized Twist oligo pool were processed to extract barcode sequences. Barcodes were extracted from positions 29-54 of read 1 and the reverse complement of positions 0-25 of read 2. Only read pairs where both extracted barcodes exactly matched known barcodes in the reference library and were identical to each other were retained for analysis. Barcode representations in the input library were calculated as the frequency of each barcode among all filtered reads.

##### Viral library sequencing and read processing

FASTQ files from paired-end sequencing of the viral supernatant library were processed to extract guide identities, clonal barcodes, and UMIs. Read 1 contained the first guide protospacer (sgRNA1), and read 2 contained both the clonal barcode and the second guide protospacer (sgRNA2). UMIs were extracted from the I1 index read. Guide protospacers were mapped to the reference library using fuzzy string matching. For each observed protospacer sequence, alignment scores were computed against all reference protospacers. Only protospacers with a minimum alignment score of 90 out of 100 were retained and assigned to the best-matching reference sequence. Clonal barcodes were required to exactly match known barcodes in the reference library. Reads were retained only if both guide sequences met the minimum alignment threshold and the clonal barcode was successfully mapped to the reference library.

##### Guide pair representation analysis

Guide pair representations in the viral library were quantified by counting the total number of reads for each combination of sgRNA1 and sgRNA2 identities among filtered reads. These counts reflect the abundance of each guide pair in the packaged viral library.

##### Barcode representation analysis

Barcode frequencies in the viral library were calculated by counting unique UMIs associated with each clonal barcode. Barcode representations in the viral library were compared to their representations in the input Twist library to assess cloning and packaging fidelity.

##### Sequencing and read processing for single-guide pilot and dual-guide genetic interaction map

FASTQ files from paired-end sequencing were processed to extract guide identities, clonal barcodes, and unique molecular identifiers (UMIs). For the single-guide pilot screen, read 1 contained the guide protospacer sequence and read 2 contained the clonal barcode. For the dual-guide screen, read 1 contained the first guide protospacer while read 2 contained both the clonal barcode and the second guide protospacer. UMIs were extracted from index reads (I1) in both screens. To prevent issues with sequencer cluster calling from constant sequences at read starts, the dual-guide screen libraries incorporated 1-7 variable bases between the Nextera Read 1 sequence and the amplification primer binding site during PCR library preparation.

Clonal barcodes, which identify cells descended from the same cell at the time of viral infection, were decoded using freebarcodes with a two-part barcode design (15bp + 10bp). The single-guide pilot screen used a reference library of 10,000 designed barcodes, and the dual-guide screen used a reference library of 120,000 designed barcodes. Guide protospacer sequences were aligned to the reference library using fuzzy string matching with a minimum similarity score of 90 out of 100 for the single-guide pilot screen and 80 out of 100 for the dual-guide screen. For the dual-guide screen, protospacer sequences were identified by fuzzy matching to the constant sequence immediately upstream of each protospacer, allowing up to 1 edit distance to account for the variable base insertions. Reads were retained only if all guide sequences and the clonal barcode were successfully mapped to the reference libraries.

For each valid read, we recorded guide identities, the mapped clonal barcode, and the UMI sequence. Reads were then grouped by guide (single-guide screen) or guide pair (dual-guide screen), clonal barcode, and sample identity. Within each group, UMIs were collapsed to count unique transcripts, generating UMI count matrices for both the identity and reporter transcripts. These UMI count matrices served as the input for downstream analysis.

##### Sequencing and read processing for single-cell genetic interaction map

FASTQ files from paired-end sequencing were processed to extract guide pair identities, clonal barcodes, partition barcodes, and unique molecular identifiers (UMIs). Read 1 contained the first guide protospacer sequence, and Read 2 contained both the clonal barcode and the second guide protospacer sequence. Index read (I1) contained partition barcodes identifying individual wells from 96-well plates, with 24 wells used per sample, and UMIs.

Clonal barcodes were decoded using freebarcodes with a two-part barcode design (15bp + 10bp) and matched against a reference library of 10,000 designed barcodes. Guide protospacer sequences were aligned to the reference library using fuzzy string matching with a minimum similarity score of 80 out of 100. Partition barcodes were aligned to the reference with a minimum similarity score of 75 out of 100. Reads were retained only if both guide sequences, the clonal barcode, and the partition barcode were successfully mapped to their respective reference libraries.

For each valid read, we recorded the identities of both guides, the mapped clonal barcode, the partition barcode, and the UMI sequence. In this screen, each cell was defined by the unique combination of guide pair, clonal barcode, and partition barcode. Reads were then grouped by guide pair, clonal barcode, partition barcode, and sample identity. Within each group, UMIs were collapsed to count unique transcripts, generating single-cell UMI count matrices for both the identity and reporter transcripts.

##### Quality control and filtering

To remove clonal lineages that could bias measurements, we filtered overrepresented lineages. For each lineage, defined by the combination of guide (or guide pair) and clonal barcode, we calculated its representation as the fraction of total UMI counts for that guide or guide pair. We then calculated a global threshold as the mean plus three standard deviations of all lineage representations across the dataset, with a minimum threshold of 5%. Overrepresented lineages exceeding this threshold for either transcript were removed from subsequent analysis. In the single-guide screen, guides were designated as controls if they were non-targeting. In the dual-guide screen, guide pairs were designated as controls if both guides in the pair were non-targeting. Replicates were processed together throughout the analysis to maximize the number of lineages per guide or guide pair, except when assessing replicate concordance for quality control purposes.

##### Compensated reporter phenotype calculation

To isolate reporter-specific signals, we performed linear regression of log2-transformed reporter counts against log2-transformed identity counts, including replicate and sample covariates to remove batch effects. This removes correlation between the two signals due to lineage size and expression bias due to integration site, and helps account for any promoter interference due to the nested design. Log2 transformation was performed with a pseudocount of 10 to stabilize variance. The regression model was fit exclusively on controls and then applied to all guides or guide pairs to predict expected reporter expression. Reporter residuals were calculated as the difference between observed and predicted log2 reporter expression.

Lineages were grouped by guide or guide pair, and the mean reporter residual was calculated for each. These mean residuals were z-score normalized using the mean and standard deviation of control guide measurements to generate reporter phenotype measurements.

##### Identity phenotype derivation

To correct for batch effects in identity transcript measurements, we used Poisson regression to model raw identity UMI counts as a function of replicate and sample covariates. The model was fit exclusively on controls and then applied to all guides or guide pairs to generate predicted identity counts. Residuals were calculated as observed minus predicted counts, then rescaled to the original UMI count range to produce batch-corrected abundance measurements for each lineage.

To quantify depletion relative to the input library, batch-corrected abundance measurements were summed by guide or guide pair to calculate total abundance in the screen. Screen representation was calculated as each guide’s or guide pair’s abundance divided by the total abundance. For the single-guide pilot screen, input library representation was determined from the cloning product. For the dual-guide screen, input library representation was determined from the packaged lentiviral library. Depletion scores were computed as the log2 fold-change between screen representation and input library representation, with a small pseudocount of 1e-7 added to avoid undefined logarithms. These depletion scores were z-score normalized using the mean and standard deviation of control measurements to generate identity phenotype measurements.

#### Single-guide pilot screen analyses

##### Statistical testing of guide-level effects

To assess the statistical significance of guide-level phenotypes, we performed Mann-Whitney U tests comparing the distribution of clonal lineage measurements for each guide against the distribution of control guide measurements. This analysis was performed separately for reporter and identity phenotypes, testing whether the distribution of measurements across clonal lineages differed significantly from controls. For each guide, we performed separate Mann-Whitney U tests against each control guide, generating multiple p-values. The median p-value across all control comparisons was used as the final test statistic for that guide. P-values were adjusted for multiple testing using the Benjamini-Hochberg false discovery rate procedure. Effect sizes were quantified using the rank-biserial correlation coefficient, with the median value across control comparisons reported for each guide.

Correlation with essential genes Perturb-seq and gene set enrichment analysis: To validate reporter phenotypes and identify transcriptional programs associated with reporter activity, we correlated guide-level reporter phenotypes with mean expression profiles from a prior Perturb-seq screen^60^ in K562 cells targeting essential genes with a substantial number of overlapping guides. For each guide in the pilot screen, perturbations in the essential genes screen were matched by protospacer sequence. For each gene in the Perturb-seq data, we calculated the Pearson correlation between the pilot guide’s reporter phenotype and the gene’s expression across matched perturbations.

Since the essential genes screen used two protospacers per gene, this yielded two correlation coefficients per gene, which were averaged to obtain a single correlation coefficient per gene. Genes were ranked by their correlation coefficients to create a ranked gene list.

To identify transcription factor regulatory programs associated with reporter activity, we performed gene set enrichment analysis (GSEA) using the prerank method with the ranked gene list described above. Enrichment was assessed against the MSigDB C3 transcription factor targets gene set collection (TFT legacy), where each gene set consists of genes whose promoters contain conserved instances of a transcription factor binding motif. Gene sets containing 100-500 genes were included in the analysis. Statistical significance was determined using 100,000 permutations, and significant gene sets were identified using a family-wise error rate (FWER) threshold of 0.05.

##### Comparison with hCRISPRi-v2 fitness scores

To validate identity phenotypes as measures of cellular fitness, we compared depletion scores from our screen with fitness scores from published hCRISPRi-v2 growth screens^81^. Guides from our screen were matched to guides from the hCRISPRi-v2 library by protospacer sequence. The correlation between depletion scores and fitness scores was assessed using Pearson correlation.

#### Dual-guide genetic interaction map analyses

##### Derivation of single and dual-guide phenotypes

From reporter and identity phenotype measurements, we derived single guide phenotypes and dual-guide phenotypes for genetic interaction analysis. Single guide phenotypes were obtained from guide pairs where one guide was a non-targeting control. For each targeting guide, we identified all measurements where that guide was paired with any control guide, averaged across both orientations (control-targeting and targeting-control) and across all control guides to obtain single guide phenotype estimates. Dual-guide phenotypes were obtained from pairs of two targeting guides. Measurements for reciprocal guide pairs were averaged when both orientations were available.

##### Genetic interaction calculation

Genetic interactions were calculated using a per-gene quadratic regression approach previously developed for fitness genetic interaction mapping^41^. For each query guide, we modeled the expected dual-guide knockout phenotype as a quadratic function of the partner guide’s single phenotype:

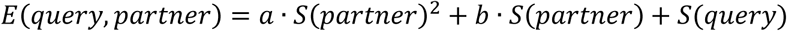

where *S(partner)* and *S(query)* are the single phenotypes of the partner and query guides, and the intercept is constrained to equal the query guide’s single phenotype.

To ensure biological plausibility, we imposed a monotonicity constraint requiring that *dE*/*dS(partner)* ≥ 0 across the range of observed single phenotypes. This constraint prevents biologically implausible scenarios where a query gene would reverse the rank ordering of phenotypic effects across all partner genes. Optimization was performed using sequential least squares programming with Huber loss (ε = 1.35) and L2 regularization (α = 0.0001) to ensure robustness against outliers.

Genetic interaction scores were calculated as the residuals between observed and expected dual-guide knockout phenotypes. For each guide pair, interaction scores from both orientations were averaged to generate symmetric genetic interaction matrices. Finally, interaction scores were scaled by dividing by the standard deviation across all pairwise interactions. This analysis was performed separately on reporter and identity phenotype measurements to generate genetic interaction maps for each phenotype.

##### Hierarchical clustering

Genetic interaction matrices were hierarchically clustered using average linkage clustering. Similarity between interaction profiles was calculated using cosine similarity in a NaN-aware manner, and distance was defined as one minus the cosine similarity. Optimal leaf ordering was applied to the dendrogram to minimize the distance between adjacent leaves.

##### Gene cluster recall

To assess which known complexes and transcriptional clusters could be recalled from the genetic interaction map, we performed neighborhood enrichment analysis using gene sets for complexes and transcriptional clusters represented in the library (by construction). For each guide in a known cluster, we calculated the fraction of neighboring guides whose targeted genes also belong to the same cluster within a sliding window. The neighborhood window extended 10 positions on each side of the query guide in the hierarchical clustering ordering, and each guide was counted independently when multiple guides targeted the same gene.

Statistical significance was assessed by permutation testing. For each permutation, guides were randomly reordered while preserving the guide identities, and neighborhood enrichment scores were recalculated. P-values were computed as the fraction of permutations with enrichment scores equal to or greater than the observed score, using 1000 permutations. P-values were corrected for multiple testing across all gene sets using the Benjamini-Hochberg false discovery rate procedure, with a significance threshold of 0.05.

For guides belonging to multiple significant clusters, we assigned each guide to a single cluster based on cluster size, enrichment scores, and proximity in the dendrogram. Among clusters within a maximum position distance of 80, assignment prioritized the largest cluster, with ties broken by enrichment scores. This approach ensures that guides are assigned to well-represented clusters while respecting their position in the ordering.

Significant guides were expanded to include neighboring guides within the same window whose targeted genes belong to the same cluster. This captures cluster members positioned near significant guides that may be at the border of the enriched region. Orphaned neighbors without nearby significant guides in the same cluster were subsequently removed. Clusters with a minimum of 2 guides after filtering were retained. The final set of significant guides and their assigned clusters represent the recalled gene sets from the hierarchical clustering.

##### Manual annotations

In addition to the automated gene cluster recall, we manually annotated guides that displayed clear clustering patterns but were not identified by the statistical enrichment analysis. Manual annotations were made based on visual inspection of the hierarchical clustering and included: (1) multiple guides targeting the same gene that clustered together, indicating reproducibility, and (2) small groups of functionally related genes (e.g., known binding partners or complex members) that clustered together but did not reach statistical significance in the enrichment analysis. These manual annotations supplemented the recalled gene sets and were visualized alongside the automated clusters in subsequent analyses.

##### Split heatmap visualization

Genetic interaction matrices were visualized using split heatmaps where the upper triangle displays genetic interaction scores scaled to unit standard deviation and the lower triangle displays dual-guide phenotypes scaled to unit standard deviation. Single-guide phenotypes were displayed in a sidebar using the same scaling as the dual-guide phenotypes.

Guides were ordered according to the hierarchical clustering dendrogram. Recalled gene clusters and manual annotations were indicated by colored bars along the matrix axes.

##### Cluster-level interaction

To summarize genetic interactions between gene clusters, we calculated average genetic interaction scores between all pairs of guides from different clusters. For each cluster pair, the mean genetic interaction score across all guide pairs was computed to generate a cluster-level interaction matrix. This analysis was performed both for recalled clusters only and for recalled clusters combined with manual annotations.

##### UMAP network visualization

To visualize the relationships between gene clusters, we projected genes onto UMAP embeddings from a prior genome-wide Perturb-seq study in K562 cells^60^. Genes were colored by their single-guide reporter phenotypes. Recalled cluster labels were positioned at the density-weighted centroid of their member genes’ UMAP coordinates. Edges between recalled clusters were drawn when their average genetic interaction score exceeded the 97.5th percentile (positive interactions) or fell below the 2.5th percentile (negative interactions) of all cluster-cluster interaction scores. Manual annotations were displayed but not included in network edge calculations. For visual clarity, each cluster was connected to a maximum of 3 other clusters, prioritized by interaction strength. Edge width was scaled proportionally to interaction strength, and edges were drawn as curved arcs to avoid overlapping with gene positions.

##### Testing recall of downsampled genetic interaction maps using √PCP

To perform downsampling analysis, all lineages from the dual-guide genetic interaction map were randomly split into two equal halves to create pseudo-replicates. For each pseudo-replicate, we subsampled varying numbers of lineages, averaging 5, 10, 15, 20, or 30 lineages per guide pair, and performed the analysis pipeline to calculate genetic interactions. This subsampling was repeated with 5 different random seeds for each lineage number.

For each subsampled dataset, genetic interactions were calculated using a global quadratic regression model rather than the per-gene approach used in the standard analysis, as sparse data were insufficient for reliable per-gene fitting. Genetic interaction scores were calculated both with and without square-root Principal Component Pursuit (√PCP) recovery. √PCP was implemented in Python following the ADMM algorithm described by Zhang et al.^78^, with modifications to handle missing values. Missing entries were masked and excluded from the data fidelity term ||*L* + *S* − *D*||_*F*_, with optimization constraints only enforced on observed entries. We used the theoretically motivated default hyperparameters λ = 1/√n_1_ and μ = √(n_2_/2). The reproducibility between pseudo-replicates was quantified by calculating the Pearson correlation coefficient across all guide pairs. Additionally, we compared genetic interactions from the subsampled dataset with an average of 5 lineages per guide pair to the full dataset (∼35 lineages) from the other pseudo-replicate to assess signal recovery.

#### Single-cell genetic interaction map analyses

##### Barcode collision rate estimation and single-cell resolution

The lentiviral library paired 10,000 distinguishable sgRNA pairs with 10,000 distinguishable clonal barcodes, yielding a theoretical diversity of *N* = 10^8^ unique vector identities. We performed two independent replicates, each transducing approximately *n* = 2.3 × 10^6^ cells. Following population expansion, cells were distributed across 24 wells per replicate for unique indexing PCR, recovering approximately 10^7^ cells per replicate as unique combinations of well index, sgRNA, and clonal barcode.

Collisions arise from two independent sources. (1) Lineage level collisions. Drawing *n* transduction events from a pool of *N* yields an expected lineage collision rate (probability that a founding cell shares its vector identity with at least one other founder) of approximately 1 – *e* 2.3% per replicate. (2) Indexing collisions introduced by cells from the same lineage that sort into the same well. With ∼10^7^ recovered cells from 2.3 × 10^6^ founders per replicate, mean lineage size is approximately 4.35 cells. For a cell to be uniquely resolved, all other members of its lineage must sort into different wells. This probability is (23/24)^n−1^ = (23/24)^3.35^ ≈ 0.87, yielding an indexing collision rate of ∼13%. This will dominate over lineage collisions because of the small number of cells per lineage. Consequently, we estimate ∼86–87% of detected cells are uniquely resolved.

These collision rates can be independently controlled by simple modifications to experimental design. First, increasing the clonal barcode pool from 10,000 to the 120,000 library used in our large genetic interaction map would reduce the lineage collision rate from ∼2.3% to ∼0.2%. Second, indexing collisions can be controlled by increasing the number of partitions. For example, performing the library prep in 96 well format would reduce the indexing collision rate from 13% to 3.4%.

##### Pseudobulk aggregation and genetic interaction calculation

To analyze genetic interactions from the single-cell genetic interaction map, single-cell measurements were aggregated into pseudobulk profiles. UMI counts were summed across all wells for each combination of guide pair, clonal barcode, sample, and well column. Well column was defined by the plate column position. The pseudobulk data were then analyzed using the standard dual-guide screen pipeline, with replicate and well column included as covariates in the regression models for reporter and identity phenotype derivation. Genetic interactions were calculated using a global quadratic regression model rather than the per-gene approach, as the relatively small scale of this screen limited the data available for per-gene fitting. Gene clusters in the genetic interaction map were manually annotated based on visual inspection of the hierarchical clustering. Annotations were assigned to genes belonging to known protein complexes or functional pathways that clustered adjacently or nearly adjacently (within 1 position) in the hierarchical ordering. These manual annotations were used to label the genetic interaction map heatmap.

##### Single-cell distribution visualization

To visualize the distribution of single-cell measurements within clonal lineages for specific perturbations, lineages were filtered to retain only those with a minimum of 2 cells. For each perturbation, lineages were ranked by their mean reporter expression, and the top 20% of lineages with the highest mean expression were selected for visualization. Log2-transformed reporter UMI counts with a pseudocount of 1 were displayed as boxplots showing the distribution within each lineage, with individual cells overlaid as points.

