## Supplemental Figures for "Scaling perturbations: beyond genome-scale CRISPR screens"

**Figure S1. Schematic of T7-mediated linear amplification of reporter and identity transcripts and flow cytometry gating strategy.**

a. Schematic of the T7-mediated linear amplification strategy used to process PORTAL reporter and identity transcripts. Because the identity transcript is fully contained within the reporter transcript, conventional PCR amplification can generate chimeric molecules that scramble the 3' ends carrying the perturbation and clonal barcode through recombination. PORTAL therefore uses a linear amplification approach in which a template-switching reverse transcription reaction installs a T7 promoter at the 5' end of each transcript and a unique molecular identifier (UMI) at the 3' end to enable single-molecule counting. Transcripts are then amplified by in vitro transcription and converted back to cDNA in a second reverse transcription step. The resulting cDNAs are produced in large quantities and differ in size, allowing physical separation of reporter and identity libraries. Finally, the 3' ends of each library containing the perturbation, clonal barcode, and UMI are independently amplified by PCR, converted into sequencing libraries, and sequenced to produce reporter and identity transcript counts.

b. Flow cytometry gating strategy for Fig. 1. Gating was as follows: cells (FSC-A, SSC-A), single-cells (FSC-A, FSC-H), sgRNA expression (GFP+), histogram of mCherry expression.

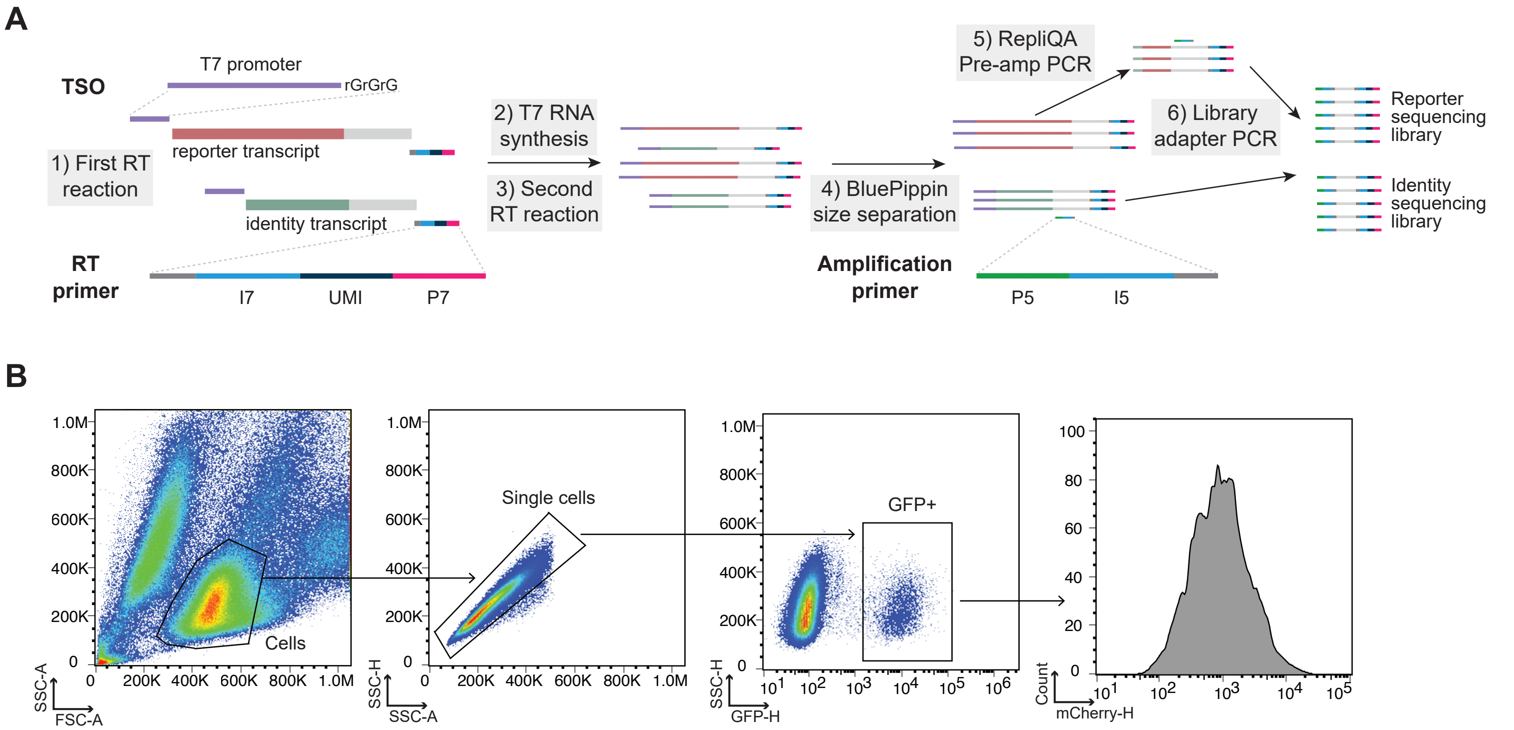
A
TSO
T7 promoter
rGrGrG
reporter transcript
1) First RT reaction
identity transcript
RT primer
I7
UMI
P7
2) T7 RNA synthesis
3) Second RT reaction
4) BluePippin size separation
Amplification primer
P5
I5
5) RepliQA Pre-amp PCR
6) Library adapter PCR
Reporter sequencing library
Identity sequencing library
B
Cells
Single cells
GFP+
SSC-A
FSC-A
SSC-H
SSC-A
SSC-H
GFP-H
Count
mCherry-H

**Figure S2. Analysis pipeline and quality control of pilot PORTAL screen.**

a. Pipeline for computing compensated reporter activity. Raw reporter and identity UMI counts show correlated variation due to differences in lineage size, total expression across integration sites, and promoter interference arising from the nested vector architecture. To account for these effects, reporter UMI counts are regressed against identity UMI counts across all measured lineages. The residuals from this regression define the compensated reporter activity. For each perturbation, this yields a lineage-resolved distribution of compensated reporter activities, which is compared to the distribution from cells containing non-targeting sgRNAs using a Mann-Whitney test to produce guide effect sizes.
b. Pipeline for computing fold depletion as a proxy for fitness. The identity transcript is constitutively expressed within each lineage and serves as a measure of lineage abundance. Identity UMI counts are first corrected for technical effects across replicates or batches using Poisson regression. Corrected counts are then summed for each sgRNA and normalized by the total across the experiment to estimate relative representation at the endpoint. Comparison of these values to sgRNA representation in the input lentiviral library yields an inferred measure of fitness-associated depletion or enrichment.
c. Validation of inferred fitness measures. Scatter plot comparing sgRNA-level fitness estimates from the pilot PORTAL screen, computed as in (b), to fitness measurements for overlapping sgRNAs in an independent K562 CRISPR screen, showing concordance between the two assays.
d. Reproducibility of sgRNA-level effect size estimates across replicates. Scatter plot comparing Mann-Whitney effect sizes for compensated reporter activity and identity across the two biological replicates of the pilot PORTAL screen.
e. Top ten enriched transcription factor signatures identified by gene set enrichment analysis. PORTAL AP-1 reporter effects were first aggregated from sgRNAs to gene-level perturbations and matched to the corresponding gene perturbations in an independent Perturb-seq dataset. Treating AP-1 reporter activity as an additional quantitative phenotype, we correlated it with transcriptome-wide gene expression changes across perturbations in the Perturb-seq data. Genes were ranked by this correlation and used as input to gene set enrichment analysis, revealing strong enrichment of AP-1-related transcription factor signatures.

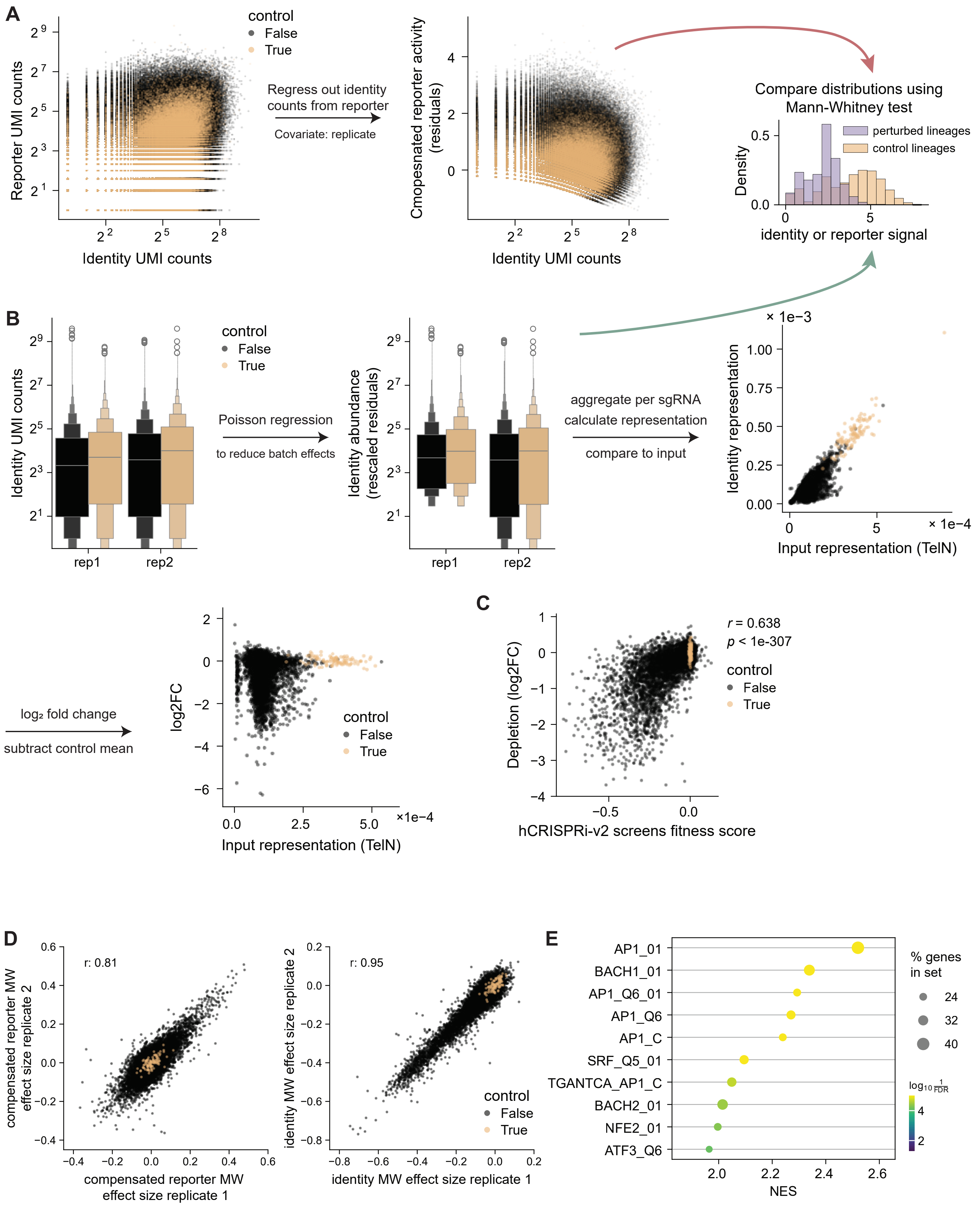
A
control
False
True
Reporter UMI counts
Identity UMI counts
Regress out identity counts from reporter
Covariate: replicate
Cmopesnated reporter activity (residuals)
Identity UMI counts
Compare distributions using Mann-Whitney test
perturbed lineages
control lineages
Density
identity or reporter signal
B
Identity UMI counts
rep1
rep2
Poisson regression
to reduce batch effects
Identity abundance (rescaled residuals)
aggregate per sgRNA
calculate representation
compare to input
Identity representation
Input representation (TelN)
log2 fold change
subtract control mean
log2FC
Input representation (TelN)
C
r = 0.638
p < 1e-307
Depletion (log2FC)
hCRISPRi-v2 screens fitness score
D
r: 0.81
compensated reporter MW effect size replicate 2
compensated reporter MW effect size replicate 1
r: 0.95
identity MW effect size replicate 2
identity MW effect size replicate 1
E
AP1_01
BACH1_01
AP1_Q6_01
AP1_Q6
AP1_C
SRF_Q5_01
TGANTCA_AP1_C
BACH2_01
NFE2_01
ATF3_Q6
NES
% genes in set
24
32
40
log10 1/FDR

**Figure S3. Quality control comparisons for CAP cloning.**

a. Long-read sequencing validation of CAP-cloned and conventionally cloned libraries. The top row reproduces the data from Fig. 3B for CAP-cloned products generated by in vitro amplification and TelN processing. The bottom row shows the corresponding analysis for libraries cloned using a conventional plasmid-based workflow, in which the assembled library was transformed into bacteria. Plots show percent sequence identity across the length of the PORTAL vector, with variable regions corresponding to the two sgRNA protospacers and the clonal barcode.
b. Structural variant rates in CAP-cloned and conventionally-cloned plasmid libraries. For the visualization in (a), molecules containing structural variants were excluded. The plot shows the frequency of structural variants detected by long-read sequencing in CAP-cloned libraries compared to traditionally cloned plasmid libraries.

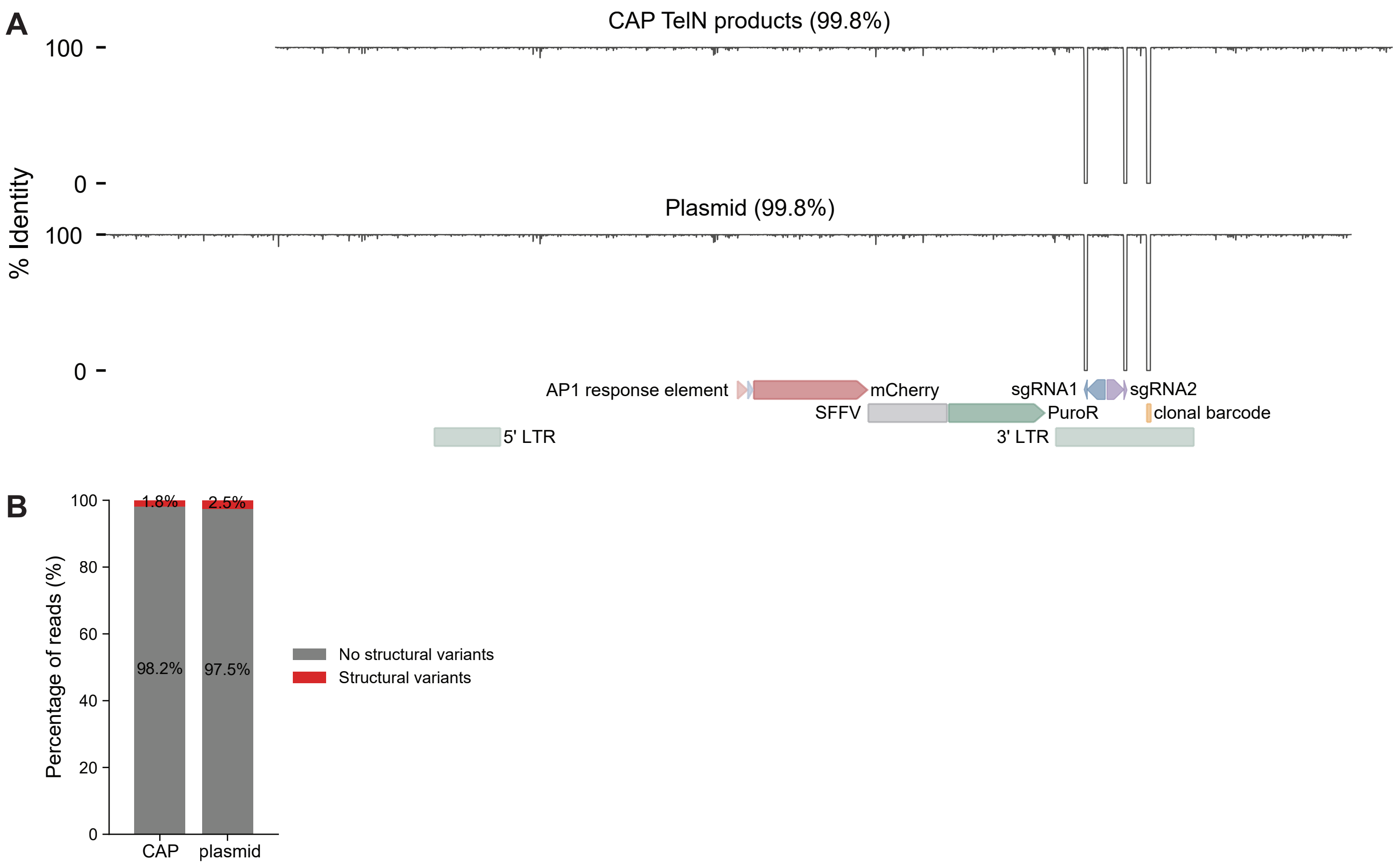
A
CAP TelN products (99.8%)
Plasmid (99.8%)
% Identity
100
0
AP1 response element
mCherry
sgRNA1
sgRNA2
SFFV
PuroR
clonal barcode
5' LTR
3' LTR
B
Percentage of reads (%)
1.8%
2.5%
98.2%
97.5%
CAP
plasmid
No structural variants
Structural variants

**Figure S4. Analysis and quality control for PORTAL AP-1 genetic interaction map.**

a. Number of clonal lineages detected per sgRNA pair in PORTAL GI map.
b. Dual-sgRNA library design. Each sgRNA constant region was cloned into both sgRNA positions and measured in both orientations, enabling assessment of technical reproducibility by comparison across orientations. PR denotes protospacer region of guide specifying the target.
c. Reproducibility of reporter and identity phenotypes across sgRNA orientations. Scatter plots compare AP-1 reporter counts (left) and identity transcript counts (right) for sgRNA pairs measured in opposite orientations. Heatmaps show conserved structure in the raw measurements.
d. Derivation of lineage-resolved reporter phenotypes for GI map, following the pipeline shown in Fig. S2A. Raw reporter and identity UMI counts show correlated variation due to differences in total expression across integration sites and promoter interference arising from the nested vector architecture (left). To account for these effects, reporter UMI counts are regressed against identity UMI counts across all measured lineages, along with replicate and sample covariates, and the residuals define the compensated reporter activity (middle). For each sgRNA pair, this yields a lineage-resolved distribution of compensated reporter activities, which are averaged across lineages. These guide-pair-level phenotypes are then $z$-normalized using the mean and standard deviation of control pairs and used for downstream genetic interaction analysis (right).
e. Framework for calculating genetic interaction (GI) scores from dual-sgRNA phenotypes. Top, schematic of dual-sgRNA library design and definition of phenotypes. For each sgRNA, pair phenotypes are measured when combined with a query sgRNA ("sgRNA pair phenotype") and compared to the phenotype of that sgRNA paired with a non-targeting guide ("sgRNA single phenotype"). Bottom left, for each query sgRNA, a guide-specific quadratic model is fit relating sgRNA pair phenotypes as a function of sgRNA single phenotypes; deviations from this expectation define positive (higher-than-expected) or negative (lower-than-expected) genetic interactions. Bottom right, GI scores are then assembled into a matrix $A$, symmetrized by averaging measurements across orientations ($A$ + $A^T$)/2, and used for clustering and downstream analysis.

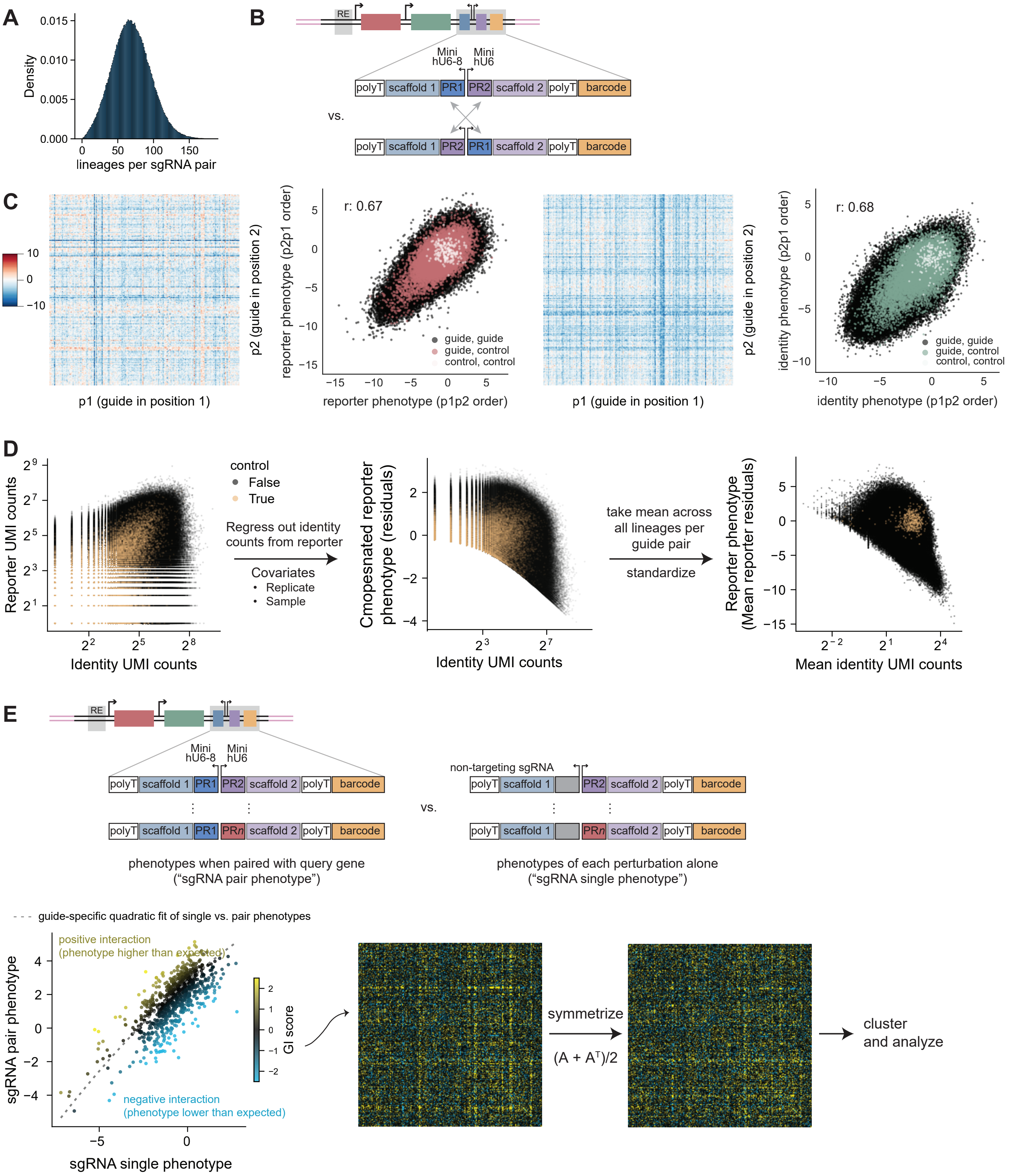
A
Density
0.015
0.010
0.005
0.000
0
50
100
150
lineages per sgRNA pair
B
RE
Mini hU6-8
Mini hU6
polyT
scaffold 1
PR1
PR2
scaffold 2
polyT
barcode
vs.
polyT
scaffold 1
PR2
PR1
scaffold 2
polyT
barcode
C
10
0
−10
p2 (guide in position 2)
p1 (guide in position 1)
r: 0.67
reporter phenotype (p2p1 order)
reporter phenotype (p1p2 order)
guide, guide
guide, control
control, control
p2 (guide in position 2)
p1 (guide in position 1)
r: 0.68
identity phenotype (p2p1 order)
identity phenotype (p1p2 order)
guide, guide
guide, control
control, control
D
Reporter UMI counts
Identity UMI counts
control
False
True
Regress out identity counts from reporter
Covariates
Replicate
Sample
Cmopesnated reporter phenotype (residuals)
Identity UMI counts
take mean across all lineages per guide pair
standardize
Reporter phenotype (Mean reporter residuals)
Mean identity UMI counts
E
RE
Mini hU6-8
Mini hU6
polyT
scaffold 1
PR1
PR2
scaffold 2
polyT
barcode
polyT
scaffold 1
PR1
PRn
scaffold 2
polyT
barcode
vs.
non-targeting sgRNA
polyT
scaffold 1
PR2
scaffold 2
polyT
barcode
polyT
scaffold 1
PRn
scaffold 2
polyT
barcode
phenotypes when paired with query gene (“sgRNA pair phenotype”)
phenotypes of each perturbation alone (“sgRNA single phenotype”)
guide-specific quadratic fit of single vs. pair phenotypes
positive interaction (phenotype higher than expected)
negative interaction (phenotype lower than expected)
sgRNA pair phenotype
sgRNA single phenotype
GI score
symmetrize
(A + A^T)/2
cluster and analyze

**Figure S5. A fitness genetic interaction map measured using PORTAL.**

a. Pipeline for computing fitness phenotypes and GI scores from PORTAL identity transcript data. Processing mirrors the fold-depletion pipeline shown in Fig. S2B, extended to dual-sgRNA measurements. Left, identity UMI counts for individual clonal lineages across replicates are corrected for batch effects (replicate and sample) using Poisson regression. Corrected identity counts are then aggregated across lineages for each sgRNA pair and compared to sgRNA-pair representation in the input lentiviral library to compute $\log_2$ fold changes in representation, serving as a proxy for fitness effects. These fitness phenotypes are z-normalized using the mean and standard deviation of control pairs. As for reporter-based GIs, guide-specific quadratic models are fit relating single-sgRNA fitness phenotypes to sgRNA-pair phenotypes, and deviations from expectation define positive and negative fitness genetic interactions.
b. Fitness genetic interaction (GI) map measured using PORTAL identity transcript counts. The lower diagonal shows inferred fitness phenotypes ($\log_2$ fold change in representation) for each sgRNA pair, while the upper diagonal shows corresponding fitness GI scores, which are used for clustering. The first strips along the bottom and right indicate fitness effects for individual gene perturbations. The second strip annotates recalled biological features, including protein complexes and genes that clustered together in Perturb-seq. Fitness phenotypes and GI scores are scaled to the standard deviation across the dataset for visualization on the same scale. While derived from the same underlying perturbations as the AP-1 reporter GI map (Fig. 4C), interaction structure is distinct.

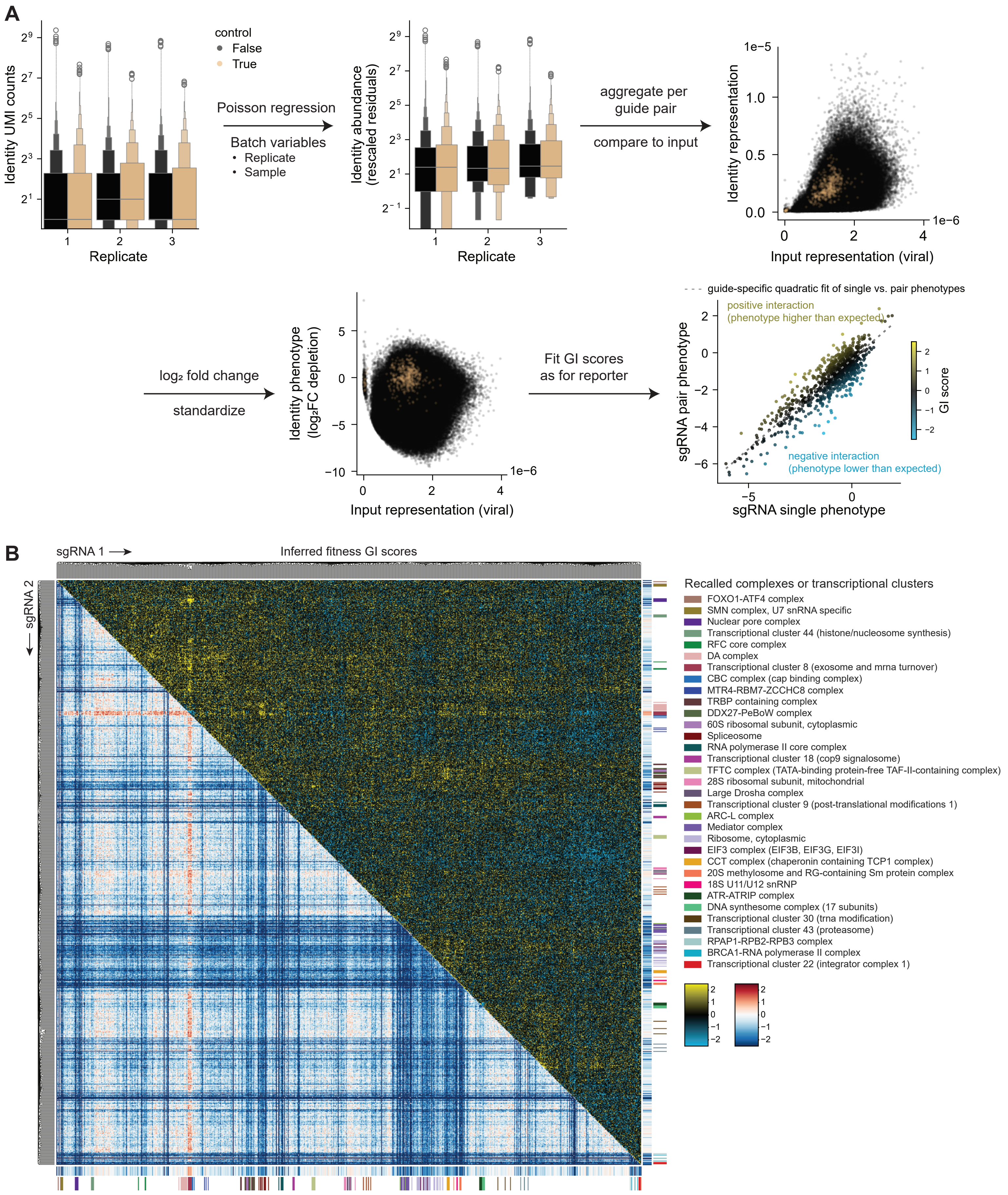
A
control
False
True
Identity UMI counts
Replicate
Poisson regression
Batch variables
• Replicate
• Sample
Identity abundance (rescaled residuals)
aggregate per guide pair
compare to input
Identity representation
Input representation (viral)
1e−5
1e−6
log₂ fold change
standardize
Identity phenotype (log₂FC depletion)
Fit GI scores as for reporter
guide-specific quadratic fit of single vs. pair phenotypes
positive interaction (phenotype higher than expected)
negative interaction (phenotype lower than expected)
sgRNA pair phenotype
sgRNA single phenotype
GI score
B
sgRNA 1
sgRNA 2
Inferred fitness GI scores
Recalled complexes or transcriptional clusters
FOXO1-ATF4 complex
SMN complex, U7 snRNA specific
Nuclear pore complex
Transcriptional cluster 44 (histone/nucleosome synthesis)
RFC core complex
DA complex
Transcriptional cluster 8 (exosome and mrna turnover)
CBC complex (cap binding complex)
MTR4-RBM7-ZCCHC8 complex
TRBP containing complex
DDX27-PeBoW complex
60S ribosomal subunit, cytoplasmic
Spliceosome
RNA polymerase II core complex
Transcriptional cluster 18 (cop9 signalosome)
TFTC complex (TATA-binding protein-free TAF-II-containing complex)
28S ribosomal subunit, mitochondrial
Large Drosha complex
Transcriptional cluster 9 (post-translational modifications 1)
ARC-L complex
Mediator complex
Ribosome, cytoplasmic
EIF3 complex (EIF3B, EIF3G, EIF3I)
CCT complex (chaperonin containing TCP1 complex)
20S methylosome and RG-containing Sm protein complex
18S U11/U12 snRNP
ATR-ATRIP complex
DNA synthesome complex (17 subunits)
Transcriptional cluster 30 (trna modification)
Transcriptional cluster 43 (proteasome)
RPAP1-RPB2-RPB3 complex
BRCA1-RNA polymerase II complex
Transcriptional cluster 22 (integrator complex 1)

**Figure S6. Pathway-level AP-1 GI map.**

a. A pathway-level GI map was constructed by averaging sgRNA-level GI scores across genes belonging to biological features recalled in the AP-1 GI map, including protein complexes, pathways, and transcriptional clusters defined in an independent Perturb-seq dataset. Only features for which multiple sgRNAs clustered together in the AP-1 GI map are shown.
b. Visualization of pathway-level GIs. Dots represent genes from the Perturb-seq dataset used to design the sgRNA library; colored dots indicate genes included in the GI map, with color reflecting AP-1 reporter activity upon knockdown. Labels indicate recalled features, with gray denoting manual annotations. Links show strong interactions between features, colored by interaction strength.

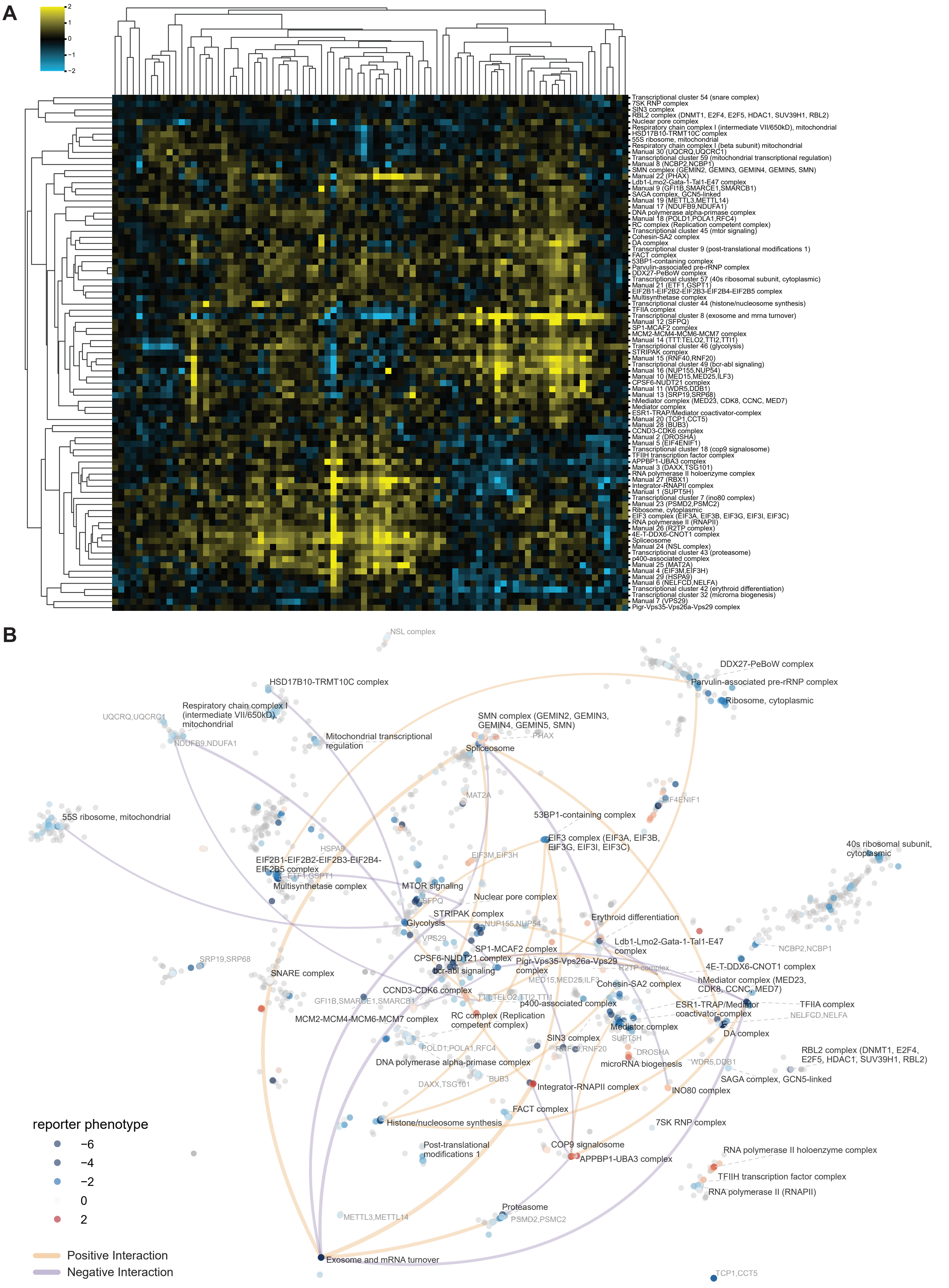
A
2
1
0
−1
−2
Transcriptional cluster 54 (snare complex)
7SK RNP complex
SIN3 complex
RBL2 complex (DNMT1, E2F4, E2F5, HDAC1, SUV39H1, RBL2)
Nuclear pore complex
Respiratory chain complex I (intermediate VII/650kD), mitochondrial
HSD17B10-TRMT10C complex
55S ribosome, mitochondrial
Respiratory chain complex I (beta subunit) mitochondrial
Manual 30 (UQCRQ,UQCRC1)
Transcriptional cluster 59 (mitochondrial transcriptional regulation)
Manual 8 (NCBP2,NCBP1)
SMN complex (GEMIN2, GEMIN3, GEMIN4, GEMIN5, SMN)
Manual 22 (PHAX)
Ldb1-Lmo2-Gata-1-Tal1-E47 complex
Manual 9 (GFI1B,SMARCE1,SMARCB1)
SAGA complex, GCN5-linked
Manual 19 (METTL3,METTL14)
Manual 17 (NDUFB9,NDUFA1)
DNA polymerase alpha-primase complex
Manual 18 (POLD1,POLA1,RFC4)
RC complex (Replication competent complex)
Transcriptional cluster 45 (mtor signaling)
Cohesin-SA2 complex
DA complex
Transcriptional cluster 9 (post-translational modifications 1)
FACT complex
53BP1-containing complex
Parvulin-associated pre-rRNP complex
DDX27-PeBoW complex
Transcriptional cluster 57 (40s ribosomal subunit, cytoplasmic)
Manual 21 (ETF1,GSPT1)
EIF2B1-EIF2B2-EIF2B3-EIF2B4-EIF2B5 complex
Multisynthetase complex
Transcriptional cluster 44 (histone/nucleosome synthesis)
TFIIA complex
Transcriptional cluster 8 (exosome and mrna turnover)
Manual 12 (SFPQ)
SP1-MCAF2 complex
MCM2-MCM4-MCM6-MCM7 complex
Manual 14 (TTT:TELO2,TTI2,TTI1)
Transcriptional cluster 46 (glycolysis)
STRIPAK complex
Manual 15 (RNF40,RNF20)
Transcriptional cluster 49 (bcr-abl signaling)
Manual 16 (NUP155,NUP54)
Manual 10 (MED15,MED25,ILF3)
CPSF6-NUDT21 complex
Manual 11 (WDR5,DDB1)
Manual 13 (SRP19,SRP68)
hMediator complex (MED23, CDK8, CCNC, MED7)
Mediator complex
ESR1-TRAP/Mediator coactivator-complex
Manual 20 (TCP1,CCT5)
Manual 28 (BUB3)
CCND3-CDK6 complex
Manual 2 (DROSHA)
Manual 5 (EIF4ENIF1)
Transcriptional cluster 18 (cop9 signalosome)
TFIIH transcription factor complex
APPBP1-UBA3 complex
Manual 3 (DAXX,TSG101)
RNA polymerase II holoenzyme complex
Manual 27 (RBX1)
Integrator-RNAPII complex
Manual 1 (SUPT5H)
Transcriptional cluster 7 (ino80 complex)
Manual 23 (PSMD2,PSMC2)
Ribosome, cytoplasmic
EIF3 complex (EIF3A, EIF3B, EIF3G, EIF3I, EIF3C)
RNA polymerase II (RNAPII)
Manual 26 (R2TP complex)
4E-T-DDX6-CNOT1 complex
Spliceosome
Manual 24 (NSL complex)
Transcriptional cluster 43 (proteasome)
p400-associated complex
Manual 25 (MAT2A)
Manual 4 (EIF3M,EIF3H)
Manual 29 (HSPA9)
Manual 6 (NELFCD,NELFA)
Transcriptional cluster 42 (erythroid differentiation)
Transcriptional cluster 32 (microrna biogenesis)
Manual 7 (VPS29)
Pigr-Vps35-Vps26a-Vps29 complex
B
NSL complex
DDX27-PeBoW complex
Parvulin-associated pre-rRNP complex
Ribosome, cytoplasmic
HSD17B10-TRMT10C complex
Respiratory chain complex I (intermediate VII/650kD), mitochondrial
UQCRQ,UQCRC1
NDUFB9,NDUFA1
SMN complex (GEMIN2, GEMIN3, GEMIN4, GEMIN5, SMN)
PHAX
Mitochondrial transcriptional regulation
Spliceosome
MAT2A
EIF4ENIF1
55S ribosome, mitochondrial
53BP1-containing complex
EIF3 complex (EIF3A, EIF3B, EIF3G, EIF3I, EIF3C)
HSPA9
EIF3M,EIF3H
40s ribosomal subunit, cytoplasmic
EIF2B1-EIF2B2-EIF2B3-EIF2B4-EIF2B5 complex
ETF1,GSPT1
Multisynthetase complex
MTOR signaling
SFPQ
Nuclear pore complex
STRIPAK complex
Glycolysis
NUP155,NUP54
Erythroid differentiation
VPS29
Ldb1-Lmo2-Gata-1-Tal1-E47 complex
SP1-MCAF2 complex
NCBP2,NCBP1
SRP19,SRP68
CPSF6-NUDT21 complex
Pigr-Vps35-Vps26a-Vps29 complex
R2TP complex
4E-T-DDX6-CNOT1 complex
SNARE complex
bcr-abl signaling
MED15,MED25,ILF3
hMediator complex (MED23, CDK8, CCNC, MED7)
CCND3-CDK6 complex
Cohesin-SA2 complex
GFI1B,SMARCE1,SMARCB1
TTT:TELO2,TTI2,TTI1
p400-associated complex
ESR1-TRAP/Mediator coactivator-complex
TFIIA complex
NELFCD,NELFA
MCM2-MCM4-MCM6-MCM7 complex
RC complex (Replication competent complex)
Mediator complex
DA complex
SIN3 complex
SUPT5H
RNF40,RNF20
DROSHA
POLD1,POLA1,RFC4
RBL2 complex (DNMT1, E2F4, E2F5, HDAC1, SUV39H1, RBL2)
DNA polymerase alpha-primase complex
microRNA biogenesis
WDR5,DDB1
BUB3
SAGA complex, GCN5-linked
DAXX,TSG101
Integrator-RNAPII complex
INO80 complex
FACT complex
Histone/nucleosome synthesis
7SK RNP complex
reporter phenotype
−6
−4
−2
0
2
Post-translational modifications 1
COP9 signalosome
APPBP1-UBA3 complex
RNA polymerase II holoenzyme complex
TFIIH transcription factor complex
RNA polymerase II (RNAPII)
Proteasome
METTL3,METTL14
PSMD2,PSMC2
Positive Interaction
Negative Interaction
Exosome and mRNA turnover
TCP1,CCT5
