## Supplemental Protocols for "Scaling perturbations: beyond genome-scale CRISPR screens"

### CAP cloning single-guide cloning

### Step 1: FastDigest digestion of parent vector with BstXI and BamHI

Parent vector: pAT005

|  | **µL per reaction** |
| --- | --- |
| Parent vector (2 µg) | X |
| FastDigest BstXI (Thermo Scientific, FD1024) | 2.5 |
| FastDigest BamHI (Thermo Scientific, FD0054) | 2.5 |
| 10X FastDigest Buffer (NOT GREEN!) | 5 |
| Water | 40 - X |

Incubate at 37ºC for 1 hr.

Clean up with 0.88X ProNex Size-Selective Beads (Promega, NG2001). Elute all DNA in TE Buffer (Invitrogen, 12090015) Elute in 25% of the ProNex® Chemistry volume used. Quantify using Nanodrop.

Take 1 µL to view on 1% EX E-GEL (Invitrogen, G401001) after purification.

#### Step 2: Twist libraries amplification

The stitching primers add the sgRNA constant region (CR) as a linker between the two libraries.

Resuspend each twist library at 5 ng/µL.

1. sgRNA guide library

Input library: CRISPRi_essentials (5 ng/µL)

Forward primer: sgRNA_stitch_F

Reverse primer: sgRNA_stitch_R

Output: sgRNA-CR

2. Barcode library

Input library: barcode twist library (at 5 ng/µL)

Forward primer: barcode_stitch_F

Reverse primer: barcode_stitch_R

Output: CR-barcode

Set up the reaction on ice. Do multiple parallel amplifications (8) to reduce the effect of PCR jackpotting. Set up each as an 8.4X master mix:

|  | **Amount** | **µL per reaction** | **8.4X MM** |
| --- | --- | --- | --- |
| Input library | 2.5 ng | 0.5 | 4.2 µL |
| Forward Primer | 0.75 µL | 0.75 | 6.3 µL |
| Reverse Primer | 0.75 µL | 0.75 | 6.3 µL |
| Nuclease-free water | Up to 25 µL | 10.5 | 88.2 µL |
| 2X KAPA HiFi HotStart Ready Mix (Roche, 07958935001) | 12.5 µL | 12.5 | 105 µL |
| **Total** |  | **25** |  |

Run the following PCR programs:

Guide library

| Step | Temperature | Time | Cycles |
| --- | --- | --- | --- |
| Initial Denaturation | 95°C | 3 min | 1 |
| Denaturation | 98°C | 20 sec | 8 |
| Annealing | 63°C | 15 sec |  |
| Extension | 72°C | 15 sec |  |
| Final Extension | 72°C | 2 min | 1 |
| Hold | 4°C | ∞ |  |

Barcode library

| Step | Temperature | Time | Cycles |
| --- | --- | --- | --- |
| Initial Denaturation | 95°C | 3 min | 1 |
| Denaturation | 98°C | 20 sec | 8 |
| Annealing | 70°C | 15 sec |  |
| Extension | 72°C | 15 sec |  |
| Final Extension | 72°C | 2 min | 1 |
| Hold | 4°C | ∞ |  |

Merge the replicate reactions of each amplification. Purify using Monarch® PCR & DNA Cleanup Kit (NEB, T1030). Elute in 25 µL TE Buffer.

Take 1 µL to view on 1% EX E-GEL after purification. Expect bands around 160 bp for sgRNA-CR and 190 bp for CR-barcode.

### Step 3: Stitching PCR

In this step we use PCR stitching to merge the two libraries. In the first step, the two libraries essentially serve as “megaprimers” for each other due to the common presence of the sgRNA scaffold sequence. We then spike in external primers and do additional rounds of PCR to target correctly stitched products.

#### Part 1: stitch the overlapping libraries

Assemble an 8.4X MM:

|  | **Amount** | **8.4X MM** |
| --- | --- | --- |
| sgRNA-CR | 20 ng |  |
| CR-barcode | 20 ng |  |
| 2X KAPA HiFi HotStart Ready Mix (Roche, 07958935001) | 25 µL | 210 µL |
| Nuclease-free water | Up to 47 µL | Up to 394.8 µL |

Split into 8 47 ul reactions in a PCR strip and run in the following PCR program.

| Step | Temperature | Time | Cycles |
| --- | --- | --- | --- |
| Initial Denaturation | 95°C | 3 min | 1 |
| Denaturation | 98°C | 10 sec | 5 |
| Annealing | 65°C | 15 sec |  |
| Extension | 72°C | 10 sec |  |
| Final Extension | 72°C | 2 min | 1 |
| Hold | 4°C | ∞ |  |

Remove reactions from thermocycler and **place on ice**! (“Hot Start” has been deactivated so polymerase will be active at room temperature!)

#### Part 2: amplify stitched product

To each reaction add the following two primers:

| sgRNA_stitch_F (10 uM) | 1.5 µL |
| --- | --- |
| barcode_stitch_R (10 uM) | 1.5 µL |

Volume should now be 50 ul. Return to the thermocycler and run the following program for all 4 reactions. Start the reaction before adding the tubes to let the PCR block cool to 4ºC. Only add the reactions once the block is cold.

| Step | Temperature | Time | Cycles |
| --- | --- | --- | --- |
| Keep reaction cold | 4°C | 5 min | 1 |
| Initial Denaturation | 95°C | 3 min | 1 |
| Denaturation | 98°C | 10 sec | 5 |
| Annealing | 65°C | 15 sec |  |
| Extension | 72°C | 10 sec |  |
| Final Extension | 72°C | 2 min | 1 |
| Hold | 4°C | ∞ |  |

Merge the reactions and purify using 1X-2X(+1X) ProNex. Elute in 100 µL. Quantify using Qubit.

Take 1 µL to view on 1% EX E-GEL after purification. The final stitched product is expected to be ~250 bp. This stitched library is then directly able to be ligated into the cut backbone in a single reaction.

#### Step 4: Ligation

Assemble the master mixes on ice, adjust the number of reactions per desired amount of cloning product:

|  | **µL per reaction** | **32X MM** |  | **2.2X Control MM** |
| --- | --- | --- | --- | --- |
| Cut backbone (100 ng) | X |  |  | 2.2X |
| Stitched library (9.8 ng) | Y |  |  | 0 |
| NEB HiFi master mix (E2621) | 10 | 320 |  | 22 |
| Water | 10 - X - Y |  |  | 22 – 2.2X |
| **Total** | 20 |  |  | 44 |

Incubate all reactions at 50°C for 1 hr.

#### Treatments post-ligation to reduce background:

Bsu36I cuts within the lacZ stuffer sequence in our vector (and so will cut any undigested pAT005).

Add 0.25 uL of Bsu36I (NEB, R0524S) directly to each reaction and incubate at 37 ºC for 15 min. Heat inactivate by incubating at 80 ºC for 20 min.

Combine ligations into one or two tube (up to 16 reactions per tube). Combine controls into one tube.

Purify by Monarch column. Elute in appropriate volume for “Nuclease-free water + Template DNA” in the next step.

### Step 5: Long-range PCR amplification of ligated vector

Forward primer: AT012F_pScreen_TelN_amp_spacer (10 µM)

Reverse primer: AT012R_pScreen_TelN_amp_spacer (10 µM)

#### PCR amplification with repliQa HiFi ToughMix® (Quantabio, 95200). Set up the same number of reactions as the previous step.

|  | **µL per reaction** | **32x** | **Control (2.2x)** |
| --- | --- | --- | --- |
| repliQa HiFi ToughMix (2X) | 25 | 800 | 55 |
| Forward primer | 1.5 | 48 | 3.3 |
| 10 µM Reverse primer | 1.5 | 48 | 3.3 |
| Nuclease-free water + Template DNA | 22 | 352x2 | 48.4 |
| **Total** | 50 |  |  |

Run in the following PCR program:

|  | Temperature | Duration |
| --- | --- | --- |
|  | 98°C | 30 sec |
| 35 cycles | 98°C | 10 sec |
|  | 68°C | 1 min 30 sec |
|  | 68°C | 2 min |
|  | 4°C | forever |

Take 1 µL to view on 1% EX E-GEL **BEFORE** purification. Expect a major band around 7000 bp for the correct product. There is usually a smaller band around 3500 bp for side products primed off LTRs – this band should be relatively faint. The 3500 bp product should be the major/only product in the control reactions.

Run a 0.8% agarose gel and purify the correct product around 7000 bp with QIAquick Gel Extraction Kit (Qiagen, 28704). Elute in 50 µL per column (12 columns for 30 reactions).

#### Step 6: Treatment with TelN Protelomerase (NEB, M0651)

1. Set up the following reaction in a microcentrifuge tube on ice. Calculate the number of parallel reactions based on the amount of PCR product from step 5.

|  | **µL per reaction** |
| --- | --- |
| PCR product (< 900 fmol TelN sites per 60 μl reaction: ~1.8 µg) | X |
| ThermoPol Reaction Buffer (10X) | 6 |
| TelN Protelomerase | 3 |
| Nuclease-free water | 51-X |
| **Total** | **60** |

2. Gently mix the reaction by pipetting up and down and microcentrifuge briefly.
3. Incubate at 30°C for 30 min followed by 75°C for 5 minutes.

Purify by Monarch column (16 columns for 45 reactions). Elute in 20 µL per column.

Expected yield for this step is around 40%.

Measure the concentration using a Nandrop.

#### Step 7: Lentiviral production from CAP cloned product

1. On Day 0, plate 1x10^6^ Lenti-X HEK293T in a 6-well
2. Day 1: prepare CAP cloned product and packaging plasmids for transfection
   a. For one 6-well:

| **DNA** | **Amount (ng)** |
| --- | --- |
| CAP cloned product | 400 |
| psPAX2 | 366 |
| pMD2G | 183 |

*Note:* Ratio of product to packaging plasmids was optimized to our application and may vary by cargo size.

3. Mix the DNA together and top up with OPTI-MEM (Gibco, 31985062) to 35.5µL.
4. In a separate tube, add 3uL of TransIT®-LT1 Transfection Reagent (Mirus Bio, MIR2304) to 15µL of OPTI-MEM and incubate for 5 minutes.
5. Add the Transfection Reagent Mix into the DNA Mix and pipette mix 3-5x.
6. Incubate the mixture for 30 minutes at room temperature.
7. After the incubation, add the mixture dropwise around the well. Swirl gently to mix.
8. Leave the plate in the incubator for 24 hours.
9. On Day 2, replace the growth media with 2mL of BSA-DMEM (10% of 1.1g/mL BSA dissolved in PBS + 90% growth media)
10. Place in incubator for 24 hours for lentiviral production to occur.
11. On Day 3, harvest the lentiviral particles by filtering the supernatant through a 0.45µm SFCA membrane.

### CAP cloning dual-guide cloning

#### Step 1: Digestion of parent vector with Esp3I and Bsu36I

Parent vector: pRCA944 or pAT011

|  | **µL per reaction** |
| --- | --- |
| Parent vector (2 µg) | X |
| Esp3I (NEB, R0734S) | 3.5 |
| Bsu36I (NEB, R0524S) | 1.5 |
| 10X rCutSmart | 5 |
| Water | 40 - X |
| **Total** | **50** |

Incubate at 37ºC for 2 hr. Incubate at 80 ºC for 20 min to inactivate.

Clean up with 0.88X ProNex Size-Selective Beads (Promega, NG2001). Elute all DNA in TE Buffer (Invitrogen, 12090015) Elute in 25% of the ProNex® Chemistry volume used. Quantify using Nanodrop.

Take 1 µL to view on 1% EX E-GEL (Invitrogen, G401001) after purification.

#### Step 2: Twist libraries amplification

In this step, the stitching primers add the mini promoters (MP) as a linker between sgRNA1 and sgRNA2 and add constant region 2 (CR2) as a linker between sgRNA2 and barcode.

Resuspend each twist library at 5 ng/µL.

1. sgRNA1 guide library

Input library: sgRNA1 twist library (5 ng/µL)

Forward primer: AT015F (10 uM)

Reverse primer: minipromoters_stitch_R (10 uM)

Output: sgRNA1-MP

2. sgRNA2 guide library

Input library: sgRNA2 twist library (5 ng/µL)

Forward primer: minipromoters_stitch_F (10 uM)

Reverse primer: AT015R_stitching (10 uM)

Output: MP-sgRNA2-CR2

3. Barcode library

Input library: barcode twist library (5 ng/µL)

Forward primer: AT016F_stitching_barcode (10 uM)

Reverse primer: AT003R (10 uM)

Output: CR2-barcode

Set up the reactions on ice. Do parallel amplifications (8) to reduce the effect of PCR jackpotting. Set up each as an 8.4X master mix:

|  | **Amount** | **µL per reaction** | **8.4X MM** |
| --- | --- | --- | --- |
| Input library | 2.5 ng | 0.5 | 4.2 µL |
| Forward Primer | 0.75 µL | 0.75 | 6.3 µL |
| Reverse Primer | 0.75 µL | 0.75 | 6.3 µL |
| Nuclease-free water | Up to 25 µL | 10.5 | 88.2 µL |
| 2X KAPA HiFi HotStart Ready Mix (Roche, 07958935001) | 12.5 µL | 12.5 | 105 µL |
| **Total** |  | **25** |  |

Run the following PCR programs:

sgRNA1 and sgRNA2 guide libraries

| Step | Temperature | Time | Cycles |
| --- | --- | --- | --- |
| Initial Denaturation | 95°C | 3 min | 1 |
| Denaturation | 98°C | 20 sec | 8 |
| Annealing | 63°C | 15 sec |  |
| Extension | 72°C | 15 sec |  |
| Final Extension | 72°C | 2 min | 1 |
| Hold | 4°C | ∞ |  |

Barcode library

| Step | Temperature | Time | Cycles |
| --- | --- | --- | --- |
| Initial Denaturation | 95°C | 3 min | 1 |
| Denaturation | 98°C | 20 sec | 8 |
| Annealing | 70°C | 15 sec |  |
| Extension | 72°C | 15 sec |  |
| Final Extension | 72°C | 2 min | 1 |
| Hold | 4°C | ∞ |  |

Merge the replicate reactions of each amplification. Purify using Monarch® PCR & DNA Cleanup Kit (NEB, T1030). Elute in 25 µL.

Take 1 µL to view on 1% EX E-GEL after purification. Expect bands around 280 bp for sgRNA1-MP, 360 bp for MP-sgRNA2-CR2, and 200 bp for CR2-barcode.

Quantify each final library using Qubit.

#### Step 3: Stitching PCR

In this step, we use PCR stitching to merge the three libraries. In the first step, the three libraries essentially serve as “megaprimers” for each other. We then spike in external primers and do additional rounds of PCR to target correctly stitched products.

#### Part 1: stitch the overlapping libraries

Assemble a 16.8X MM:

|  | **Amount** | **16.8X MM** |
| --- | --- | --- |
| sgRNA1-MP | 20 ng |  |
| MP-sgRNA2-CR2 | 20 ng |  |
| CR2-barcode | 20 ng |  |
| 2X KAPA HiFi HotStart Ready Mix (Roche, 07958935001) | 25 µL | 420 µL |
| Nuclease-free water | Up to 47 µL | Up to 789.6 µL |

Split into 16 47 µL reactions in a PCR strip and run in the following PCR program.

| Step | Temperature | Time | Cycles |
| --- | --- | --- | --- |
| Initial Denaturation | 95°C | 3 min | 1 |
| Denaturation | 98°C | 10 sec | 5 |
| Annealing | 65°C | 15 sec |  |
| Extension | 72°C | 10 sec |  |
| Final Extension | 72°C | 2 min | 1 |
| Hold | 4°C | ∞ |  |

Remove reactions from thermocycler and **place on ice**! (“Hot Start” has been deactivated so polymerase will be active at room temperature!)

#### Part 2: amplify stitched product

To each reaction add the following two primers:

| AT015F (10 uM) | 1.5 µL |
| --- | --- |
| AT003R (10 uM) | 1.5 µL |

Volume should now be 50 µL. Return to the thermocycler and run the following program for all 4 reactions. Start the reaction before adding the tubes to let the PCR block cool to 4ºC. Only add the reactions once the block is cold.

| Step | Temperature | Time | Cycles |
| --- | --- | --- | --- |
| Keep reaction cold | 4°C | 5 min | 1 |
| Initial Denaturation | 95°C | 3 min | 1 |
| Denaturation | 98°C | 10 sec | 5 |
| Annealing | 65°C | 15 sec |  |
| Extension | 72°C | 10 sec |  |
| Final Extension | 72°C | 2 min | 1 |
| Hold | 4°C | ∞ |  |

Merge the reactions and purify using 1.25X ProNex Size-Selective Beads (Promega, NG2001). Elute in 150 µL. Quantify using Qubit.

Take 1 µL to view on 1% EX E-GEL after purification. The final stitched product is expected to be ~515 bp. This stitched library is then directly able to be ligated into the cut backbone in a single reaction.

#### Step 4: Ligation

Assemble the master mixes on ice, adjust the number of reactions per desired amount of cloning product:

|  | **µL per reaction** | **32X MM** |  | **2.2X Control MM** |
| --- | --- | --- | --- | --- |
| Cut backbone (100 ng) | X |  |  | 2.2X |
| Stitched library (33 ng) | Y |  |  | 0 |
| NEB HiFi master mix (E2621) | 10 | 320 |  | 22 |
| Water | 10 - X - Y |  |  | 22 – 2.2X |
| **Total** | 20 |  |  | 44 |

Incubate all reactions at 50°C for 1 hr.

Combine ligations into one or two tube (up to 16 reactions per tube). Combine controls into one tube.

Purify by Monarch column. Elute in appropriate volume for “Nuclease-free water + Template DNA” in the next step.

### Step 5: Long-range PCR amplification of ligated vector

Forward primer: AT012F_pScreen_TelN_amp_spacer (10 µM)

Reverse primer: AT012R_pScreen_TelN_amp_spacer (10 µM)

#### PCR amplification with repliQa HiFi ToughMix® (Quantabio, 95200). Set up the same number of reactions as the previous step.

|  | **µL per reaction** | **32x** | **Control (2.2x)** |
| --- | --- | --- | --- |
| repliQa HiFi ToughMix (2X) | 25 | 800 | 55 |
| Forward primer | 1.5 | 48 | 3.3 |
| 10 µM Reverse primer | 1.5 | 48 | 3.3 |
| Nuclease-free water + Template DNA | 22 | 352x2 | 48.4 |
| **Total** | 50 |  |  |

Run in the following PCR program:

|  | Temperature | Duration |
| --- | --- | --- |
|  | 98°C | 30 sec |
| 35 cycles | 98°C | 10 sec |
|  | 68°C | 1 min 30 sec |
|  | 68°C | 2 min |
|  | 4°C | forever |

Take 1 µL to view on 1% EX E-GEL **BEFORE** purification. Expect a major band around 7000 bp for the correct product. There is usually a smaller band around 3500 bp for side products primed off LTRs – this band should be relatively faint. The 3500 bp product should be the major/only product in the control reactions.

Run a 0.8% agarose gel and purify the correct product around 7000 bp with QIAquick Gel Extraction Kit (Qiagen, 28704). Elute in 50 µL per column (12 columns for 30 reactions).

#### Step 6: Treatment with TelN Protelomerase (NEB, M0651)

4. Set up the following reaction in a microcentrifuge tube on ice. Calculate the number of parallel reactions based on the amount of PCR product from step 5.

|  | **µL per reaction** |
| --- | --- |
| PCR product (< 900 fmol TelN sites per 60 μl reaction: ~1.8 µg) | X |
| ThermoPol Reaction Buffer (10X) | 6 |
| TelN Protelomerase | 3 |
| Nuclease-free water | 51-X |
| **Total** | **60** |

5. Gently mix the reaction by pipetting up and down and microcentrifuge briefly.
6. Incubate at 30°C for 30 min followed by 75°C for 5 minutes.

Purify by Monarch column (16 columns for 45 reactions). Elute in 20 µL per column.

Expected yield for this step is around 40%.

Measure the concentration using a Nanodrop.

#### Step 7: Lentiviral production from CAP cloned product

12. On Day 0, plate 1x10^6^ Lenti-X HEK293T in a 6-well
13. Day 1: prepare CAP cloned product and packaging plasmids for transfection
    a. For one 6-well:

| **DNA** | **Amount (ng)** |
| --- | --- |
| CAP cloned product | 400 |
| psPAX2 | 366 |
| pMD2G | 183 |

*Note:* Ratio of product to packaging plasmids was optimized to our application and may vary by cargo size.

14. Mix the DNA together and top up with OPTI-MEM (Gibco, 31985062) to 35.5µL.
15. In a separate tube, add 3uL of TransIT®-LT1 Transfection Reagent (Mirus Bio, MIR2304) to 15µL of OPTI-MEM and incubate for 5 minutes.
16. Add the Transfection Reagent Mix into the DNA Mix and pipette mix 3-5x.
17. Incubate the mixture for 30 minutes at room temperature.
18. After the incubation, add the mixture dropwise around the well. Swirl gently to mix.
19. Leave the plate in the incubator for 24 hours.
20. On Day 2, replace the growth media with 2mL of BSA-DMEM (10% of 1.1g/mL BSA dissolved in PBS + 90% growth media)
21. Place in incubator for 24 hours for lentiviral packaging.
22. On Day 3, harvest the lentiviral particles by filtering the supernatant through a 0.45µm SFCA membrane.

### Bulk library prep

#### Step 0.1: Total RNA extraction

Extract total RNA from fresh/frozen cell pellets using QIAGEN RNeasy® Midi kit (Qiagen, 75144) according to manufacturer’s protocol. Homogenization of cell lysates step: use 20G Sterile Hypodermic Needles (Air-Tite™, 14-817-208) and sterile RNase-free syringes to homogenize cell lysates by passing the lysate at least 5 times.

Add Protector RNase Inhibitor (Roche, 3335402001) and store at -80 °C if not proceeding immediately.

#### Step 0.2: mRNA isolation

Isolate mRNA using PolyATtract® mRNA Isolation Systems III (Promega, Z5300) according to manufacturer’s protocol. Elute in 100 µL of water – second elution in the protocol was not performed since we found it to only add ~2% of yield, and pooling would significantly dilute the concentration.

Add Protector RNase Inhibitor (Roche, 3335402001) and store at -80 °C if not proceeding immediately.

#### Step 1: Produce template-switched cDNA

#### In this step, reverse transcription is performed using RT primers that bind directly to the transcripts downstream of barcode with indexing and UMI.

Template-switching oligo with a T7 promoter sequence is added to introduce a T7 promoter to the 5’ end of the cDNA.

Kit: Maxima H Minus Double-Stranded cDNA Synthesis Kit (Thermo Scientific, K2562)

Set up parallel reactions for each sample on ice:

|  | **µL per reaction** |
| --- | --- |
| RT primer (100 µM): Pscreen_7XX | 2 |
| Water | 24 - X |
| RNA template (~2 µg of mRNA) | X |
| **Total** | 26 |

- Mix gently, centrifuge briefly and incubate at 65 °C for 5 min. Chill on ice.
- Add the following components in the indicated order to each reaction:

|  | **µL per reaction** |
| --- | --- |
| Template switching oligo (75 µM):  TN006-TSO_T7 | 2 |
| 4X first strand reaction mix | 10 |
| First strand enzyme mix | 2 |
| **Total** | 14 |

Total volume: 40 µL

| Temperature | Duration |
| --- | --- |
| 42°C | 90 min |
| 85°C | 5 min |
| 4°C | forever |

#### Step 2: T7 RNA synthesis

Kit: HiScribe® T7 Quick High Yield RNA Synthesis Kit (NEB, E2050L)

Assemble the same number of parallel reactions as in step 1 at room temperature in the following order:

|  | **µL per reaction** |
| --- | --- |
| NTP Buffer Mix | 50 |
| T7 RNA Polymerase Mix | 10 |
| Template DNA (pool and split) | 40 |
| **Total** | 100 |

Alternatively, when thermocycler available for larger reaction volumes (eg. PIPseq Dry bath), pool reactions for each sample and assemble at room temperature in the following order:

(Buffer Mix to Polymerase Mix ratio should remain 5:1, the amount of each component and total volume is flexible and can be adjusted according to the total reaction volume in step 1)

|  | **µL per reaction** |
| --- | --- |
| NTP Buffer Mix | 80 |
| T7 RNA Polymerase Mix | 16 |
| Template DNA (combined) | 200 |
| **Total** | 296 |

Run the following program:

| Temperature | Duration |
| --- | --- |
| 42°C | 2 hours |
| 4°C | forever |

Add 4 µL (or 10 µL for large reactions) of DNAseI and incubate at 37°C for 15 minutes.

Combine all reactions and purify the RNA using 1.8X RNAClean XP (Beckman Coulter, NC0068576). Elute in 200 µL H_2_O. Measure the RNA concentration using Nanodrop 2000.

Add Protector RNase Inhibitor (Roche, 3335402001) and store -80 °C.

#### Step 3: Second RT reaction/second strand synthesis

Kit: Maxima H Minus Double-Stranded cDNA Synthesis Kit (Thermo Scientific, K2562)

Set up parallel reactions for each sample on ice.

|  | **µL per reaction** |
| --- | --- |
| RNA template previous step (up to 25 µg) | X |
| RT primer (100 µM): P7 | 4 |
| Water | 24 - X |
| **Total** | 28 |

- Mix gently, centrifuge briefly and incubate at 65 °C for 5 min. Chill on ice.
- Add the following components in the indicated order to each reaction:

|  | **µL per reaction** |
| --- | --- |
| 4X first strand reaction mix | 10 |
| First strand enzyme mix | 2 |
| **Total** | 12 |

Total volume: 40 µL

| Temperature | Duration |
| --- | --- |
| 50°C | 45 min |
| 85°C | 5 min |
| 4°C | forever |

Combine reactions for each sample and purify using 3X ProNex Size-Selective Beads (Promega, NG2001). Elute in water in total volume for “First strand cDNA synthesis ProNex elute” across all reactions.

Set up same number of parallel reactions for second strand synthesis:

|  | **µL per reaction** |
| --- | --- |
| First strand cDNA synthesis ProNex elute | 75 |
| 5X second strand synthesis mix | 20 |
| Second strand enzyme mix | 5 |
| **Total** | 100 |

For each reaction:

- Mix gently and centrifuge briefly
- Incubate at 16 °C for 2 hr.
- Stop the reaction by adding 6 µL 0.5 M EDTA, pH 8.0 and mixing gently.
- Add 10 µL RNAseI and incubate for 5 min at RT to remove residual RNA.

Combine reactions for each sample and purify using 1.1X ProNex Size-Selective Beads (Promega, NG2001). Elute in 80 µL TE buffer (can adjust). Measure concentration using Nanodrop 2000.

#### Step 4: BluePippin to separate the two transcripts

Cassette: 0.75% DF Low Voltage 1-6kb Marker S1 (SAGE SCIENCE, BLF7510)

Selection range should be adjusted based on the expected sizes of transcripts.

Settings we used:

Single-guide GFP library

| **Target Transcript** | **Start BP** | **End BP** |
| --- | --- | --- |
| Identity | 1100 | 1680 |
| Reporter | 2200 | 3100 |

Dual-guide GFP library

| **Target Transcript** | **Start BP** | **End BP** |
| --- | --- | --- |
| Identity | 1120 | 1690 |
| Reporter | 2250 | 3150 |

Dual-guide puro library

| **Target Transcript** | **Start BP** | **End BP** |
| --- | --- | --- |
| Identity | 1003 | 1583 |
| Reporter | 2103 | 3003 |

#### Step 5: Reporter transcript-specific long-range pre-amplification

For reporter transcript only:

- Forward primer: P017seqF (10 µM)
- Reverse primer: P7 (10 µM)

Set up (8) parallel reactions using repliQa HiFi ToughMix® (Quantabio, 95200):

|  | **µL per reaction** | **8.4x** |
| --- | --- | --- |
| BluePippin DNA (+ water) | 11 | 92.4 |
| repliQa HiFi ToughMix (2X) | 12.5 | 105 |
| Forward primer | 0.75 | 6.3 |
| Reverse primer | 0.75 | 6.3 |
| **Total** | 25 |  |

- Mix gently and then centrifuge briefly to collect solutions to the bottom of tubes.
- Incubate in a thermocycler, with lid temperature set at ≥100°C, and perform PCR with the following cycling condition

|  | Temperature | Duration |
| --- | --- | --- |
|  | 98°C | 30 sec |
| 8 cycles | 98°C | 10 sec |
|  | 68°C | 45 sec |
|  | 68°C | 2 min |
|  | 4°C | forever |

Purify using 0.98X ProNex Size-Selective Beads (Promega, NG2001). Elute in 168 ul H_2_O (for 8 reactions in the next step).

#### Step 6: PCR amplification with Q5

Set up (8) parallel reactions using NEBNext® Ultra™ II Q5® Master Mix (NEB, M0544S).

*Use a different index forward primer for the two libraries (e.g. 501 for identity, 502 for reporter)

#### Identity transcript

Forward primer (10 uM):

- Single-guide library: pScreen_5XX
- Dual-guide library: pScreen_dual_5XX

Reverse primer (10 uM):

- P7

|  | **µL per reaction** | **8.4x** |
| --- | --- | --- |
| BluePippin selected template | 10 |  |
| NEBNext Ultra II Q5 Master Mix | 25 | 210 |
| Reverse primer (10 uM) | 2.5 | 21 |
| Water | 10 | 84 |
| Total | 47.5 |  |
| Well-specific | | |
| Forward primer (10 uM) | 2.5 |  |
| **Total volume** | **50** |  |

#### Reporter transcript

Forward primer (10 uM):

- Single-guide library: pScreen_5XX
- Dual-guide library: pScreen_dual_5XX

Reverse primer (10 uM):

- P7

|  | **µL per reaction** | **8.4x** |
| --- | --- | --- |
| Pre-amplified template | 20 |  |
| NEBNext Ultra II Q5 Master Mix | 25 | 210 |
| Reverse primer (10 uM) | 2.5 | 21 |
| Total | 47.5 |  |
| Well-specific | | |
| Forward primer (10 uM) | 2.5 |  |
| **Total volume** | **50** |  |

- Mix gently and then centrifuge briefly to collect solutions to the bottom of tubes.
- Incubate in a thermocycler, with lid temperature set at ≥100°C, and perform PCR with the following cycling condition

| Step | Temperature | Time | Cycles |
| --- | --- | --- | --- |
| Initial Denaturation | 98°C | 30 sec | 1 |
| Denaturation | 98°C | 10 sec | 8 cycles (identity)  6 cycles (reporter) |
| Annealing/Extension | 65°C | 2 min 30 sec |  |
| Final Extension | 65°C | 5 min | 1 |
| Hold | 4°C | ∞ |  |

Single-guide library: 1.1X-1.4X(+0.3X) ProNex to purify. Elute in 22 µL. Check on Qubit & bioanalyzer (High Sensitivity dsDNA or TapeStation).

Dual-guide library: 1X-1.3X(+0.3X) ProNex to purify. Elute in 22 µL. Check on Qubit & bioanalyzer.

Further size-selection/purification with ProNex might be needed if left-over primers observed on bioanalyzer.

Further amplification might be needed if amount of product not enough for sequencing – see optional P5/P7 amplification step on next page.

#### Step 7 (Optional): Further P5/P7 amplifications with Q5

Kit: NEBNext® Ultra™ II Q5® Master Mix (NEB, M0544S)

|  | **Volume (µL)** |
| --- | --- |
| Template | 20 |
| NEBNext Ultra II Q5 Master Mix | 25 |
| Forward primer: P5 (10 uM) | 2.5 |
| Reverse primer: P7 (10 uM) | 2.5 |
| **Total volume** | **50** |

- Mix gently and then centrifuge briefly to collect solutions to the bottom of tubes.
- Incubate in a thermocycler, with lid temperature set at ≥100°C, and perform PCR with the following cycling condition

| Step | Temperature | Time | Cycles |
| --- | --- | --- | --- |
| Initial Denaturation | 98°C | 30 sec | 1 |
| Denaturation | 98°C | 10 sec | 4 cycles |
| Annealing/Extension | 65°C | 2 min 30 sec |  |
| Final Extension | 65°C | 5 min | 1 |
| Hold | 4°C | ∞ |  |

*The number of cycles can be adjusted

Single-guide library: 1.1X-1.4X(+0.3X) ProNex to purify. Elute in 22 µL. Check on Qubit & bioanalyzer.

Dual-guide library: 1X-1.3X(+0.3X) ProNex to purify. Elute in 22 µL. Check on Qubit & bioanalyzer.

Further size-selection/purification with ProNex might be needed if left-over primers observed on bioanalyzer.

Single-guide library example bioanalyzer trace:

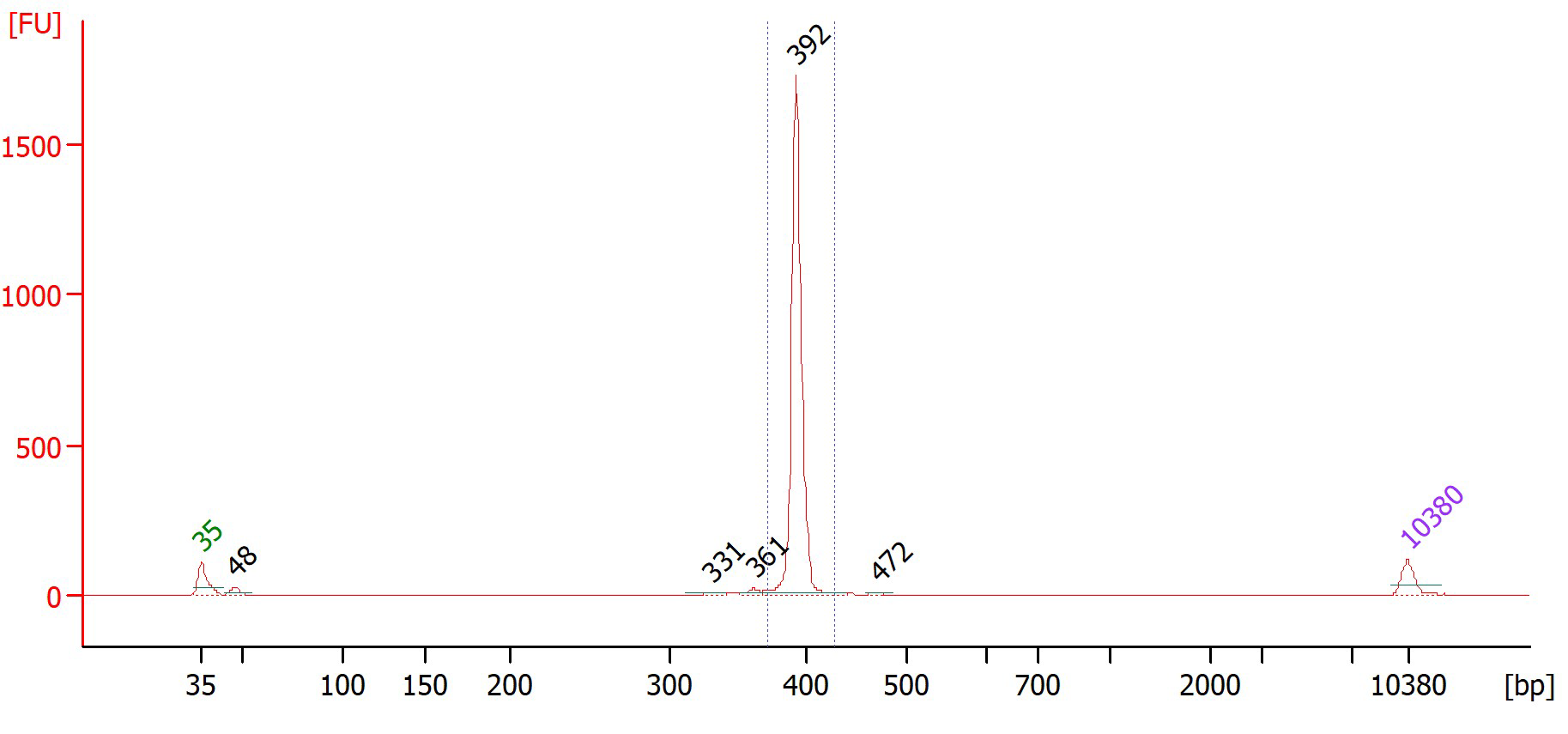

Dual-guide library example bioanalyzer trace:

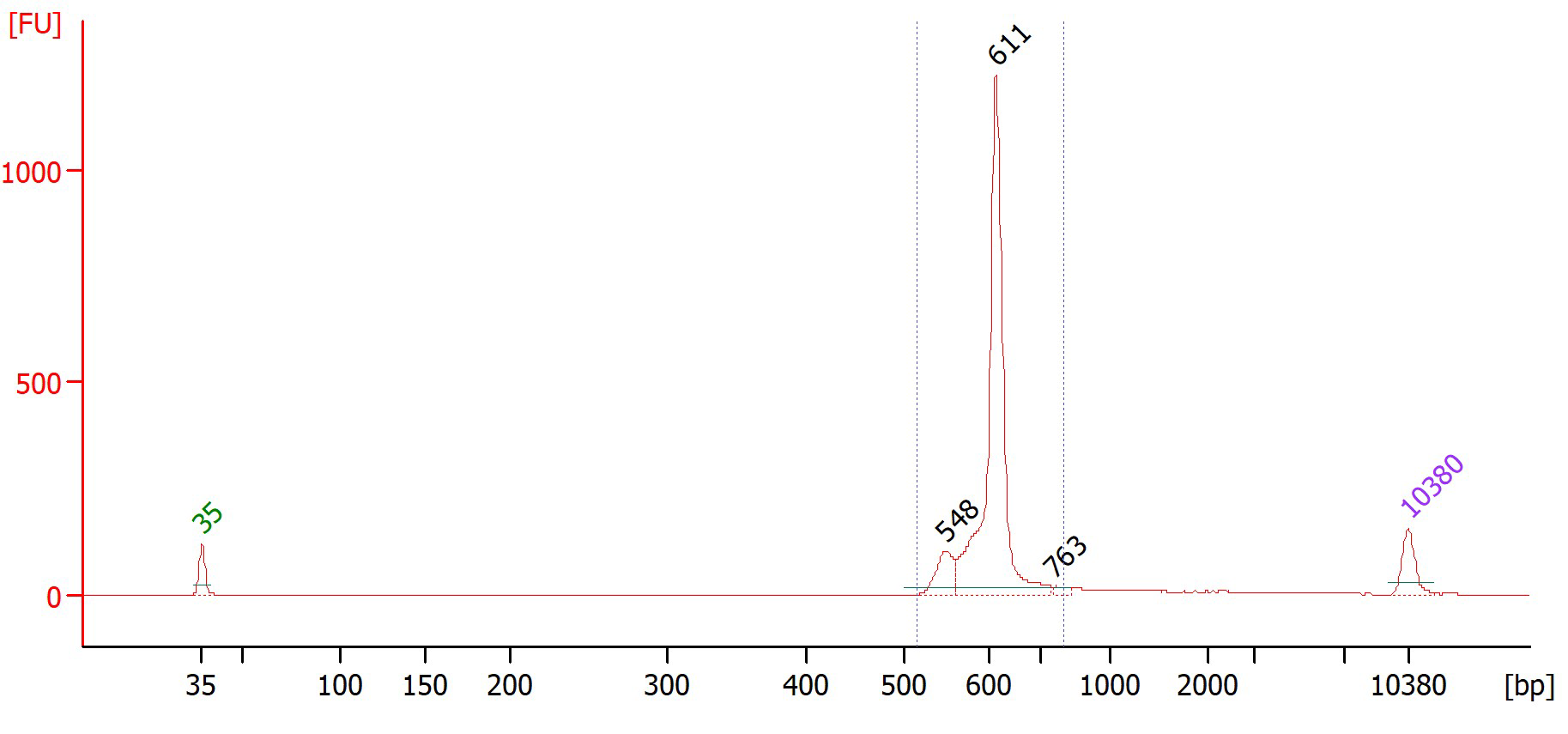

### Single-cell library prep

#### Step 0: mRNA isolation using mRNA Catcher^TM^ PLUS 96-well plate (Invitrogen, K157002)

##### Before Starting

1. Bring all reagents to room temperature
2. Mix 2X Lysis Buffer well. If the buffer contains precipitates, incubate at 37°C to solve the precipitates
3. Prepare fresh 2X Lysis Buffer with DTT by adding 25 μl 0.5 M DTT to 975 μl 2X Lysis Buffer (final DTT concentration is 5 mM) to prepare 1 ml of 1X Lysis Buffer with DTT. Mix well and use for preparing the cell lysate. Depending on the amount and type of samples you are preparing, you can scale-up the volumes accordingly.

##### Preparing lysates

Transfer suspension cells into RNase free tubes. Centrifuge to pellet cells and remove the media. Resuspend the cell pellet into an appropriate volume of PBS to obtain 8.3 x 10^6^ cells/ml.

Transfer 30 μl cell suspension containing 250,000 cells into each well of the mRNA Catcher plate. Add equal volume (30 μl) of 2X Lysis Buffer with DTT to each well. Mix well by pipetting up and down.

##### Hybridization

1. Cover the mRNA Catcher PLUS plate containing samples with adhesive foil plate cover.
2. Incubate the plate for 45-60 minutes at room temperature for RNA hybridization.

##### Washing

1. Aspirate the lysates from wells. Be sure not to scrape the well sides during aspiration.
2. Add 100 μl **Wash Buffer (W15)** to the wells.
3. Incubate the plate for 1 minute at room temperature.
4. Aspirate the Wash Buffer. Repeat washing to obtain a total of 3 washes.
5. **DO NOT REMOVE** the wash buffer after the last wash.

#### Step 1: Produce template-switched barcoded cDNA

Kit: Maxima H Minus Double-Stranded cDNA Synthesis Kit (Thermo Scientific, K2562)

#### Reverse transcription

Prepare the Elution Mix using the table below.

|  | **Volume per well (µL)** | **Master mix (µL)** |
| --- | --- | --- |
| Water | 51 |  |
| 4X first strand reaction mix | 10 |  |
| Total | 61 |  |
| Well-specific | | |
| RT primer (400 µM) | 1 |  |
| Total | 62 |  |

- Remove the Wash Buffer and immediately add 62 μl of the Elution Mix prepared as above to each well of the plate.
- Mix gently, centrifuge briefly and incubate at 65 °C for 5 min. Chill on ice.
- Add the following components in the indicated order to each well:

|  | **Volume per well (µL)** | **Master mix (µL)** |
| --- | --- | --- |
| Template switching oligo (75 µM):  TN006-TSO_T7 | 4 |  |
| 4X first strand reaction mix | 10 |  |
| First strand enzyme mix | 4 |  |
| Total | 18 |  |

Total volume: 80 µL

Mix well and cover the mRNA Catcher PLUS plate with adhesive aluminum foil plate cover.

Perform the RT reaction by incubating the plate in a thermocycler suitable for 96-well plates:

| Temperature | Duration |
| --- | --- |
| 42°C | 90 min |
| 85°C | 5 min |
| 4°C | forever |

The synthesized cDNA in solution can be stored at -20°C.

#### Step 2: T7 RNA synthesis

Kit: HiScribe® T7 Quick High Yield RNA Synthesis Kit (NEB, E2050L)

Pool cDNA per column from the plate. Assemble reactions at room temperature in the following order (one reaction per column):

|  | Volume (µL) | Added? |
| --- | --- | --- |
| Nuclease-free water | - |  |
| NTP buffer mix | 120 |  |
| T7 RNA Polymerase Mix | 24 |  |
| Template DNA | 640 |  |
| Total | 784 |  |

Incubate using the PIPseq Dry bath:

| Temperature | Duration |
| --- | --- |
| 42°C | 2 hours |
| 4°C | forever |

Add 4 µL DNAseI to each reaction and incubate at 37°C for 15 minutes.

Purify the RNA using 1.8X RNAClean XP (Beckman Coulter, NC0068576). Elute in 70 µL H_2_O per reaction. Measure the RNA concentration using Nanodrop 2000.

Add RNase inhibitor and store -80 °C.

#### Continue to step 3 through 7 in bulk library prep protocol.
